# Heterochronous laminar myelination in the human prefrontal cortex

**DOI:** 10.1101/2025.01.30.635751

**Authors:** Valerie J. Sydnor, Daniel Petrie, Shane D. McKeon, Alyssa Famalette, Will Foran, Finnegan J. Calabro, Beatriz Luna

**Affiliations:** Department of Psychiatry, University of Pittsburgh Medical Center, University of Pittsburgh, Pittsburgh, PA, USA; The Center for the Neural Basis of Cognition, University of Pittsburgh and Carnegie Mellon University, Pittsburgh, PA, USA; Department of Bioengineering, University of Pittsburgh, Pittsburgh, PA, USA

## Abstract

The human prefrontal cortex (PFC) exhibits markedly protracted developmental plasticity, yet whether reductions in plasticity occur synchronously across prefrontal cortical layers is unclear. Animal studies have shown that intracortical myelin consolidates developing circuits by restricting ongoing neuronal plasticity. Here, we use longitudinal myelin-sensitive imaging collected at ultra-high field to investigate whether superficial and deep PFC layers exhibit different timeframes of malleability. We find that myelin matures earlier in deep than in superficial compartments of the cortical ribbon; this laminar divergence in maturational timing is differentially expressed across cytoarchitecturally and functionally distinct frontal regions. By integrating myelin mapping with EEG and behavioral phenotyping, we provide evidence that prefrontal myelin impacts timescales of neural activity, task learning rates, and cognitive processing speed. Heterochronous myelination across deep and superficial layers is an underrecognized mechanism through which human association cortex balances cognitively-relevant increases in circuit stability and efficiency with extended neuroplasticity.

## Introduction

The human prefrontal cortex (PFC) is an evolutionarily-expanded core of association cortex^1–3^ that supports advanced cognitive processing and exhibits remarkably protracted development during childhood and adolescence^3–7^. The PFC’s capacity to develop over decades suggests that it balances developmental increases in circuit stability necessary for reliable cognitive functioning with continued circuit malleability fundamental to experience-dependent refinement. In this study, we investigate the hypothesis that this balance is achieved through the heterochronous maturation of a neurobiological regulator of plasticity across layers of the PFC. Far from being a uniform entity, the PFC is a mosaic of both functionally diverse cortical regions and distinct cortical layers that exhibit substantial developmental variability^3,5,7^. Prior work investigating this variability has revealed that reductions in plasticity progress hierarchically across frontal regions^5,8–11^, facilitated by asynchronous increases in intracortical myelin, a primary regulator of neuronal plasticity^5,12–14^. However, how plasticity temporally unfolds across prefrontal cortical layers—which differ in their evolutionary histories, connectivity targets, neurobiological features, and functional roles—is a fundamental unknown in our understanding of association cortex developmental chronology. Here, we harness 7 Tesla (7T), quantitative, myelin-sensitive imaging collected longitudinally to characterize temporal refinements in plasticity across deep and superficial layers of the developing frontal cortex and relate PFC myelination to neurocognitive specialization.

The prefrontal cortical ribbon has a classic laminar architecture defined by the presence of up to six cortical layers. Although layers are not equivalent across frontal regions^3,15^, they conform to a canonical circuit architecture with stereotypical input connections and output projections. Layers 1 and 4 are non-pyramidal “input” layers that receive substantial input from the thalamus and other cortical regions. Layers 5 and 6 are pyramidal layers in deep cortex that serve as the main cortical “output” layers. These layers contain pyramidal neurons that predominantly project to the subcortex, including to the thalamus, striatum, brainstem, and spinal cord, with a smaller proportion sending feed-back cortical projections^16^. Layers 2 and 3 are pyramidal layers in superficial cortex that are the main source of local and long-range cortico-cortical connections, especially feed-forward projections^16^. Layers 2/3 are proportionally thicker in the PFC of primates than in other mammalian species^17^ and have extensive local recurrent excitatory connections^16^ with high integrative capacity and coding dimensionality^18^. Layers 2/3 are thus considered the main “computation” layers of the PFC^19,20^.Despite these marked differences between cortical layers, it is not clear whether they exhibit unique maturational trajectories that differentially support early structural consolidation versus continued plasticity within the developing association cortex.

Intracortical myelin is a primary neurobiological regulator of cortical developmental plasticity^21–23^. Myelin forms in the cortex around the axons of both pyramidal neurons^24^ and parvalbumin (PV) inhibitory interneurons^25,26^: the two main neural substrates for remodeling during periods of heightened developmental plasticity^27,28^. Once myelin forms, myelin-associated inhibitors (Nogo, MAG, OMgp)^29^ interact with receptors expressed on neurons to inhibit neurite outgrowth^30^, spine turnover^21,31^, and synaptic plasticity^32^. As a result, myelin formation serves to consolidate the physical wiring of circuitry and terminate developmental windows of neuronal plasticity. Notably, although myelin formation restricts the structural plasticity of neurons, myelin itself remains relatively plastic throughout life^33–36^. Indeed, in animal models, activity-dependent myelin sheath formation and restructuring continues into adulthood and is instrumental for learning and long-term memory^34,37–40^. Intracortical myelin thus serves as both a key limiter of developmental plasticity and a structural foundation for lifelong, cognition-linked circuit adaptability.

Prior elegant studies using *in vivo* imaging to study the development of intracortical myelin have found that deep-to-mid compartments of the prefrontal cortical ribbon myelinate more than superficial compartments during the adolescent period^11,41–43^. Post-mortem studies of human and non-human primates have furthermore shown that as compared to pyramidal neurons in deep layer 5, neurons in superficial layer 3 of the adolescent PFC show more pronounced pruning of dendritic spines^6^ and slower accumulation of proteins associated with stable dendritic morphology^10^. These data convergently suggest that superficial prefrontal layers may exhibit greater plasticity than deep layers during youth; however, whether superficial and deep layers differ in the timing or duration of developmental windows of plasticity is not known. This gap in knowledge precludes an understanding of whether distinct layers of association cortex differentially contribute to the human brain’s prolonged developmental malleability and to maturation-associated changes in cortical signaling and cognitive abilities. This gap exists, in part, given that conventional MRI field strengths and contrast-based acquisitions are limited in their capacity to study refinements in plasticity regulators across deep and superficial layers with adequate sensitivity and specificity. In this study, we overcome these limitations by combining 7T MRI, quantitative neuroimaging, and intracortical depth profiling to chart the temporal maturation of myelin throughout superficial, middle, and deep compartments of the frontal cortical ribbon. Unlike contrast-based approaches that examine relative differences in signals with arbitrary units, quantitative imaging techniques provide calibrated measures of the biochemical composition of tissue expressed in physical units (e.g., proton relaxation times). Here, we specifically employ quantitative R1, a histologically-validated, myelin-sensitive imaging measure^44–47^ that increases with greater myelin density and that has excellent scan-rescan reliability at 7T^48^.

We hypothesized that developmental change in R1 would terminate earlier in deep as compared to superficial portions of the prefrontal cortical ribbon, providing evidence for earlier consolidation of layer 5/6 projections to subcortex and hierarchically lower regions together with protracted plasticity in layer 2/3 cortico-cortical connections. Furthermore, we hypothesized that laminar variability in R1 development would differ in a systematic manner across hierarchically and functionally diverse frontal regions, linking prior reports of heterogeneous PFC myelination^5,12–14^ to differences in layer-stratified myelination processes. Finally—leveraging electroencephalogram (EEG) and cognitive assessments collected from the same participants—we studied the impact of superficial and deep frontal cortex myelin on the maturation of circuit electrophysiology and PFC-dependent cognitive processes. The formation of myelin around excitatory pyramidal cells and inhibitory PV interneurons supports fast, high frequency interactions between excitatory (E) and inhibitory (I) firing^49^ and increases the speed and fidelity of ongoing neural activity^50,51^. We therefore hypothesized that higher R1-indexed myelin would be associated with faster dynamical changes in E/I population activity as well as with more efficient learning and processing speed. Altogether, we uncover heterochronous myelin maturation across deep and superficial layers of the PFC that allows higher-order association cortex to co-express stability and malleability at different stages of layer-stratified processing hierarchies.

## Results

### Quantitative imaging of myelin in the cortical ribbon

We studied spatiotemporal variability in the maturation of myelin throughout the frontal cortical ribbon in an accelerated longitudinal sample of 140 individuals ages 10-32 years old with 1-3 imaging timepoints each. Myelin was measured with quantitative R1 data (the longitudinal relaxation rate in sec^-1^) collected with 1 mm isotropic resolution at ultra-high field (7T). To characterize laminar variability in myelin density, we used intracortical profiling^11,41,43^ to quantify R1 at a range of cortical depths between pial and white matter boundaries. Specifically, we mapped volumetric R1 data to individual-specific cortical surfaces in 10% thickness increments between 0% of cortical thickness (pial boundary) and 100% of cortical thickness (gray-white boundary) (**Fig. 1a**). We then further analyzed 7 intracortical depths where >90% of the R1 signal was estimated to come from cortical gray matter (**Fig. 1b**). These 7 depths spanned 20% to 80% of cortical thickness and thus ranged approximately from the bottom of layer 2 to above mid-layer 6^19,43^ (**Fig. 1c**).

**Figure 1.**
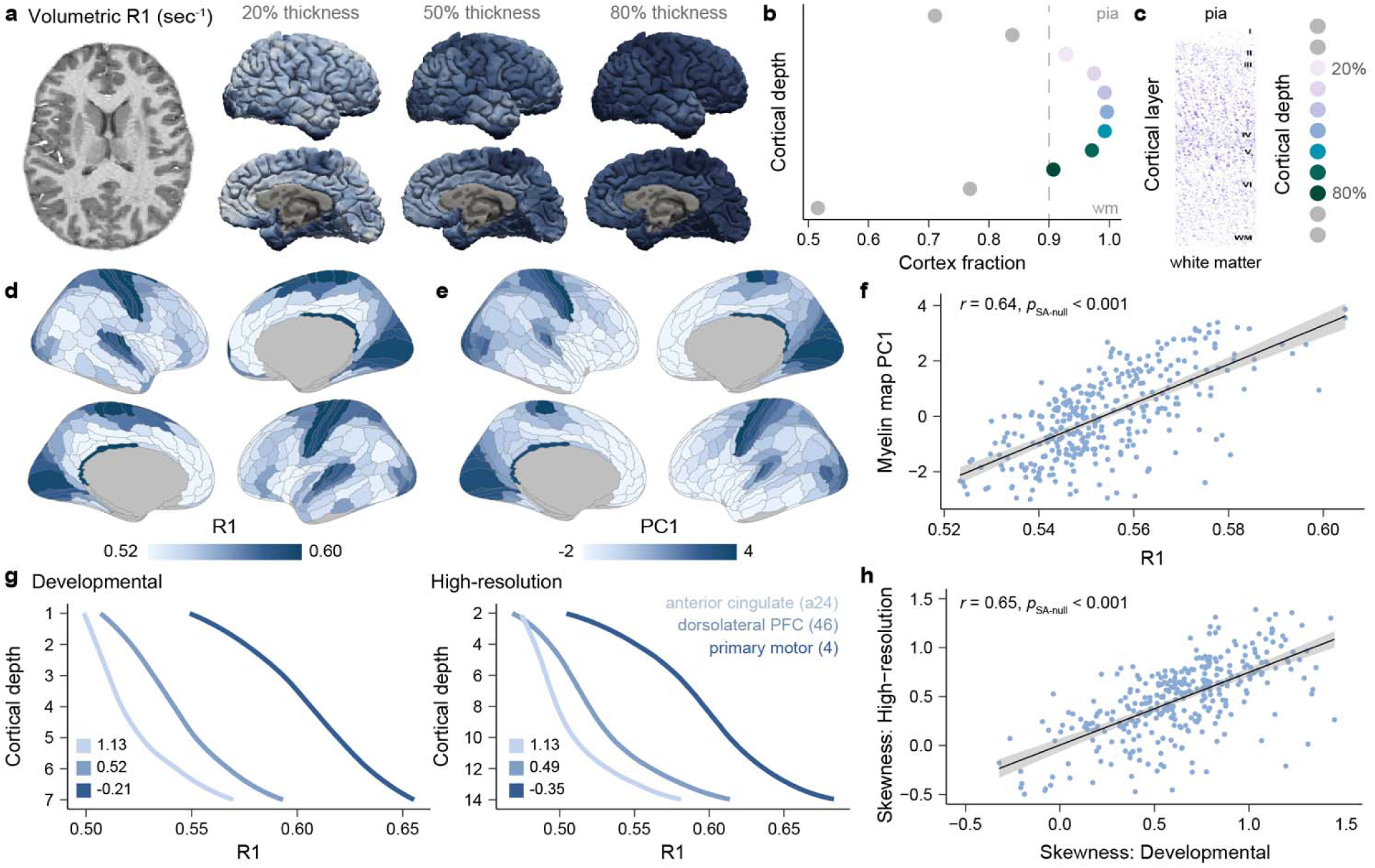
R1-based myelin mapping in the cortical ribbon. **a**) A volumetric R1 map from a representative individual is shown along with projections of R1 to that individual’s cortical surface at 20%, 50%, and 80% of cortical thickness. R1 is a quantitative MRI measure with units of sec^-1^. **b**) The intracortical depth profiling approach samples data in 10% thickness increments between 0% of cortical thickness (pial boundary) and 100% of cortical thickness (white matter boundary). To ensure that R1 was studied at depths where the signal nearly exclusively originated from gray matter, the cortex volume fraction was estimated at each cortical depth. 7 depths with an average cortex volume fraction > 0.9 were further analyzed. **c**) To provide intuition into the sampling of R1 across intracortical depths, a histochemical myelin stain of the dorsolateral PFC from García-Cabezas et al.^15^ is included, along with the original labeling of cortical layers 1-6 and white matter. The estimated locations of the 7 cortical depths under study, which range from 20% to 80% of total cortical thickness, are indicated. **d**) A map of regional average R1 in the 7T developmental dataset. **e**) The principal component (PC1) of spatial variation in three independent myelin-sensitive measures is shown. PC1 was derived from previously published cortical maps of the T1w/T2w ratio (data from Baum et al.^12^), magnetization transfer saturation (data from Hettwer et al.^41^), and myelin basic protein gene expression (data from Wagstly et al.^52^). **f**) R1 and PC1 are strongly spatially correlated across cortical regions, confirming correspondence between R1 and independent myelin mapping techniques. **g**) The shape of depth-dependent R1 profiles systematically differs between three laminarly distinct regions in the developmental dataset and a high-resolution dataset collected with 0.5 mm isotropic voxels. Numerical values in the lower left of each plot indicate the skewness of the R1 profile for the anterior cingulate (area a24; light blue), the dorsolateral prefrontal cortex (area 46; medium blue), and the primary motor cortex (area 4; dark blue). **h**) Regional variability in the skewness of depth-dependent R1 profiles is highly correlated across the cortex between the developmental and high-resolution datasets.

Prior to using these data to study prefrontal myelin development, we set out to establish that R1—as assessed by our 7T acquisition in a developmental sample—captured the expected distribution of myelin across both cortical regions and intracortical depths. We first sought to confirm that the regional distribution of R1 aligned with myelination patterns observed using other myelin mapping techniques. To accomplish this, we averaged R1 throughout the cortical ribbon within each cortical region (**Fig. 1d**) and surveyed whether regional differences aligned with the principal component of spatial variation in three independent and commonly employed myelin-sensitive measures: the T1w/T2w ratio^12^, magnetization transfer saturation^41^, and myelin basic protein gene expression^52^. The first principal component explained 76% of the variance in these three measures (PC1; **Fig. 1e**) and exhibited a robust spatial correlation to cortical R1 in our sample (*r* = 0.64, *p*_SA-null_, < 0.001; **Fig. 1f**).

We next aimed to confirm that our 1mm isotropic R1 data could capture the depth-dependent organization of myelin across cortical layers. Cross-species studies have shown that myelin increases between layers 2 and 6, with the laminar profile of increase varying between cortical regions with different degrees of laminar differentiation^11,22,47,53,54^. This variation in laminar myelination manifests as regional differences in the skewness (asymmetry) of depth-dependent myelin profiles^11^. To validate that our developmental data had adequate resolution to capture laminar shifts in myelin, we thus compared the shape and skewness of regional depth-dependent R1 profiles to those derived in high-resolution 7T R1 data collected with 0.5 mm isotropic voxels. In both the developmental dataset and the high-resolution dataset, the shape of depth-dependent R1 profiles differed between cortical regions with distinct laminar architectures (**Fig. 1g**). Furthermore, regional variability in the skewness of depth-dependent R1 profiles was highly correlated across the cortex between the developmental dataset and the high-resolution dataset (*r* = 0.65, *p*_SA-null_, < 0.001; **Fig. 1h**). Together, these findings confirm that our developmental R1 data capture key features of myeloarchitecture in the cortical ribbon—including features observable in independent myelin mapping techniques and sub-millimeter R1 data.

### Studying myelin in superficial, middle, and deep frontal cortex

A primary goal of this work was to test the hypothesis that myelin maturational trajectories differ between superficial layers 2/3 and deep layers 5/6 within the developing PFC. Having now confirmed that our developmental R1 data captured the expected regional and depth-dependent distribution of myelin, we next endeavored to ensure that we could reliably distinguish R1 signal between superficial and deep cortex throughout the frontal lobe. We first calculated the thickness of frontal lobe regions for all participants in the sample and found that regional cortical thickness predominantly ranged from 2.0 to 3.0 mm in participants of all ages (**Fig. 2a**; average frontal thickness = 2.6 mm ± 0.2 across regions). These cortical thickness values suggested that all of the frontal cortical ribbon contained R1 signal from 2-3 or more voxels—and thus 2 or 3 distinct cortical compartments. Indeed, 98.1% of cortical surface vertices in the frontal lobe contained R1 signal from 2 or more unique voxel compartments, with 72.7% of vertices containing R1 signal from 3 or more voxel compartments (**Fig. 2b**). As a result—in every single participant regardless of age—the most frequent number of independent compartments observed within the frontal cortex was 3 (**Fig. 2c**).

**Figure 2.**
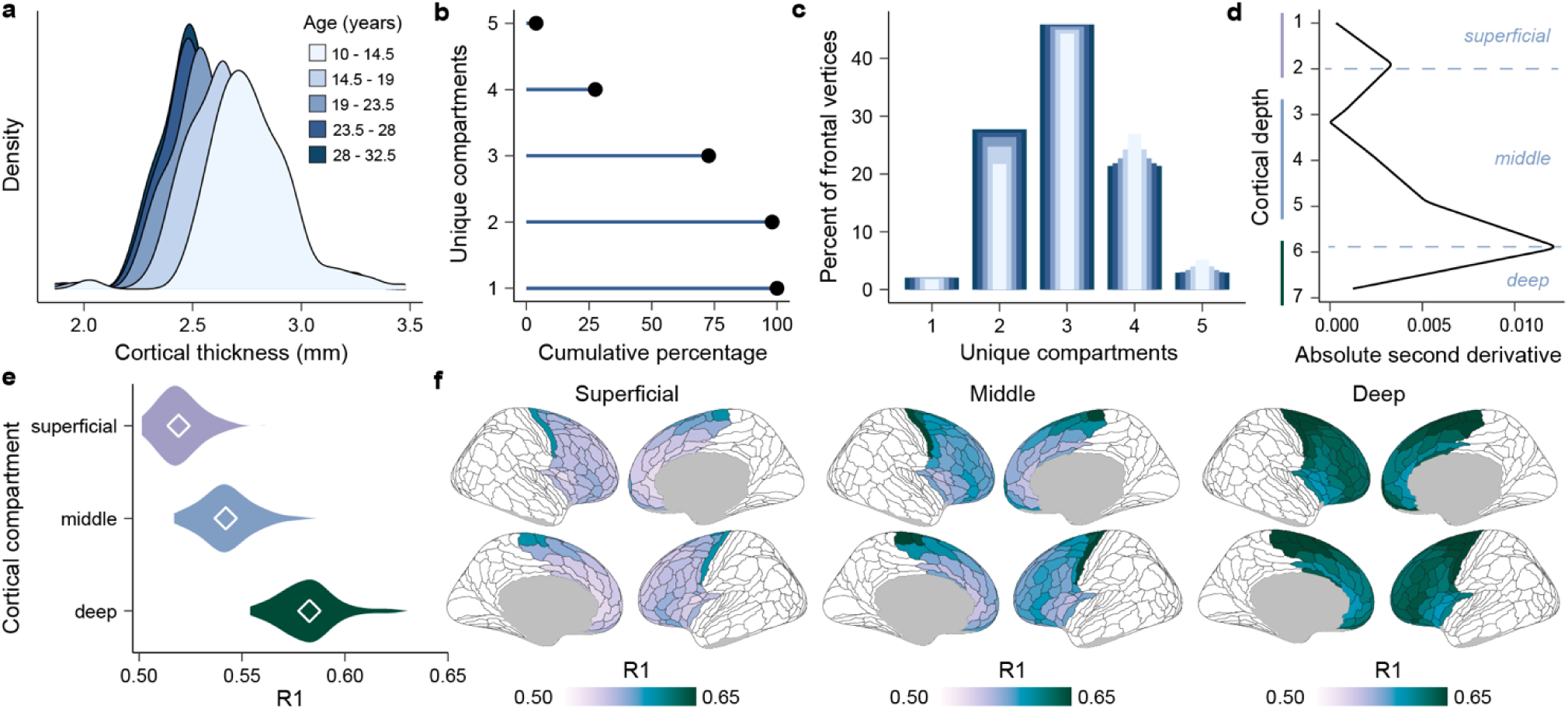
Differentiating R1 in superficial, middle, and deep compartments of the frontal cortex. **a**) A plot displaying the probability density of cortical thickness values across all frontal lobe regions is shown, stratified into 5 age groups. The thickness of all frontal lobe regions ranges from approximately 2.0 to 3.5 mm in individuals of all ages. **b**) The cumulative percentage of frontal cortex vertices that have at least N unique voxel compartments within the cortical ribbon is plotted. 27.5% of vertices contained data from 4 or more unique voxels within the cortical ribbon. 72.7% of vertices contained data from 3 or more unique voxels. 98.1% of vertices contained data from 2 or more unique voxels. **c**) The percent of frontal lobe vertices with exactly 1, 2, 3, 4, or 5 unique voxels present within the cortical ribbon is indicated. Most locations in the frontal cortical ribbon contain independent R1 signal from 3 or more voxels, allowing for the differentiation of 3 unique cortical compartments. **d**) The absolute second derivative of depth-dependent R1 change in the frontal lobe is maximal near depths 2 and 6 in the developmental data, designating data-driven boundaries between superficial, middle, and deep cortical compartments based on transitions in the statistical properties of laminar R1 profiles. **e**) R1 increases between superficial, middle, and deep compartments of the frontal cortical ribbon. Violin plots summarize the distribution of frontal region R1 within superficial, middle, and deep cortex; white diamonds represent the average frontal R1 in each cortical compartment. **f**) Cortical maps depict regional R1 values in superficial, middle, and deep cortical compartments for the frontal lobe.

These data substantiate that R1 can be reliably differentiated in superficial, middle, and deep cortex across the frontal lobe, enabling the study of how myelin matures independently in relatively more superficial layers, deep layers, and a middle transition zone.

To determine how to most appositely divide the depth-dependent frontal R1 data into superficial, middle, and deep cortical compartments, we identified where along the frontal cortical ribbon the statistical properties of R1 changed, suggestive of layer-related changes in myelination^55^. For this analysis, we leveraged the high-resolution (0.5 mm isotropic) R1 data for its enhanced ability to differentiate between cortical layers and translated findings to our developmental dataset. To directly facilitate this translation, we calculated frontal lobe R1 at 14 intracortical depths in the 0.5 mm high-resolution data, corresponding to the 7 analyzed intracortical depths in our 1 mm developmental data. We then computed the absolute second derivative of R1 across frontal intracortical depths in both datasets to identify transition points where the rate of R1 change abruptly shifted. A high absolute second derivative indicates that the depth-dependent change in R1 sharply accelerates or decelerates, as expected near boundaries between laminarly distinct cortical compartments.

In the high-resolution dataset, two peaks in the absolute second derivative were observed between cortical depths 3-4 and 11-12, uncovering data-driven boundaries between superficial, middle, and deep cortical compartments; the same peaks were observed whether R1 was averaged across the frontal lobe or examined within individual frontal regions with different degrees of laminar differentiation (**Supplementary Fig. 2.1**). These results suggested that the optimal corresponding compartment boundaries in the developmental dataset should exist near depths 2 and 6. In line with this prediction, analysis of the second derivative in the developmental sample identified data-driven boundaries at depths 2 and 6 (**Fig. 2d**). We therefore averaged R1 across depths 1-2, depths 3-5, and depths 6-7—deriving an overall measure of myelin in superficial, middle, and deep portions of the frontal cortical ribbon (**Fig. 2e, f**) that we could study developmentally.

### Heterochronous laminar myelination in the frontal cortex

To chart developmental change in R1 in superficial, middle, and deep frontal cortex, we began by averaging R1 across all frontal regions within each cortical compartment and fitting compartment-specific generalized additive mixed models (GAMMs) with age splines. R1 significantly increased with age in the frontal lobe in superficial, middle, and deep compartments (*p*_FDR_ < 0.001 in all 3 models).

However, developmental splines showed variability in the magnitude of R1 increase and the shape of age-related curves across cortical compartments (**Fig. 3a; Supplementary Fig. 3.1**). In particular, R1 exhibited shallower and more continuous increases in superficial frontal cortex. With increasing distance into the cortex, R1 showed larger age-related increases that slowed over time, resulting in trajectories that plateaued in young adulthood. An interaction model that tested whether age splines were significantly different across the 3 cortical compartments was significant (*F* = 12.15; *p* < 0.001). The same overarching pattern emerged when studying age-related change in R1 across the original 7 intracortical depths, confirming that averaging R1 within compartments did not bias developmental trajectories (**Supplementary Fig. 3.2**). In a complementary set of analyses, we calculated within-individual change in superficial, middle, and deep compartment R1 between longitudinal imaging timepoints, thereby deriving person-specific developmental trajectories. Within-person change in frontal cortex R1 was positive in >80% of individuals in all 3 compartments, increased in magnitude with greater distance into the cortex, and decelerated in late adolescence (**Supplementary Fig. 3.3**). These analyses reveal robust longitudinal increases in R1 at the individual level and confirm convergence between group-level GAMMs and within-person developmental effects.

**Figure 3.**
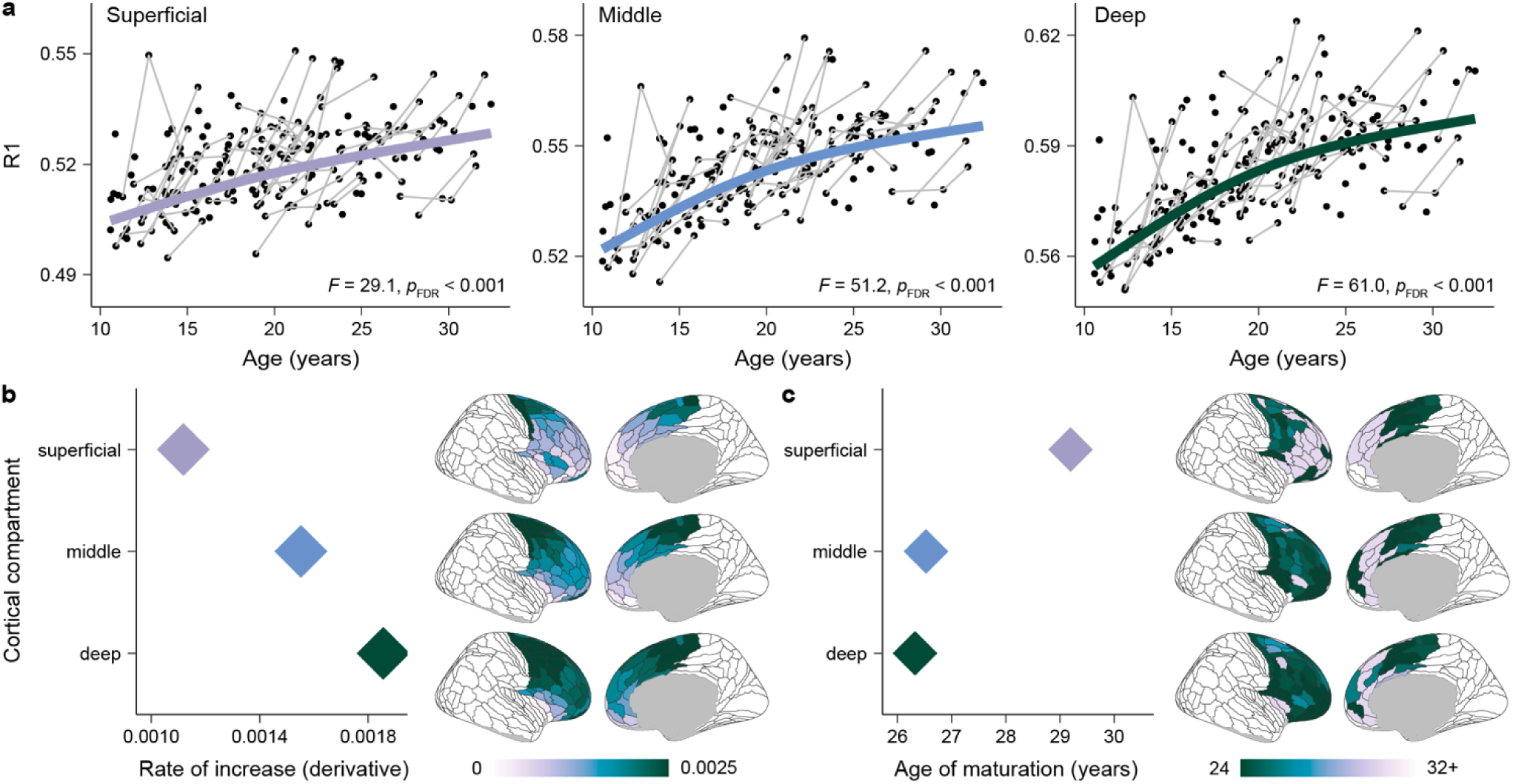
R1 matures heterochronously in superficial and deep frontal cortex. **a**) Developmental trajectories of R1, modeled using GAMM smooth functions, are shown for superficial, middle, and deep compartments of the frontal cortical ribbon. Developmental trajectories are overlaid on participant-level data; longitudinal imaging sessions are connected. The *F*-statistic and the FDR-corrected *p*-value of the age smooth term from each GAMM are indicated; *F-*statistics progressively increase in magnitude from superficial to deep cortex. **b**) The average rate at which R1 increased with age gets faster when moving from superficial to deep compartments, indicative of larger magnitude developmental change in deeper cortex. Cortical plots (right) display the average rate of R1 increase in each frontal region within superficial, middle, and deep cortex. The corresponding plot (left) quantifies the average rate of R1 increase across all frontal regions in superficial, middle, and deep cortex. The rate of increase was calculated as the average first derivative of the age spline. **c**) The age at which R1 matured in each region gets younger when moving from superficial to deep compartments, suggestive of earlier maturational timing in deeper cortex. Cortical plots (right) show the age of R1 maturation in each frontal region within superficial, middle, and deep cortex. The corresponding plot (left) quantifies the average age of maturation across frontal regions in each compartment. The age of R1 maturation was operationalized as the youngest age at which the first derivative of the age spline was no longer significantly different from zero, based on a 95% simultaneous interval.

Given initial evidence for asynchronous myelin maturation within the cortical ribbon of the entire frontal lobe, we extended our investigation to individual frontal regions by fitting compartment-specific GAMMs in each region. Nearly all frontal lobe regions showed significant increases in R1 (*p*_FDR_ < 0.05) in all 3 compartments; the percent of significant regional effects was 90% in superficial cortex, 94% in the middle transition zone, and 99% in deep cortex. By quantifying region-specific rates of R1 increase (**Fig. 3b**) and region-specific ages of R1 maturation (**Fig. 3c**), we discovered a clear developmental pattern: when moving from superficial to deep cortex, the rate of R1 developmental change increased, yet the age of R1 maturation decreased. Hence, with increasing laminar depth, age-related increases in R1 were larger in magnitude and were instantiated at a faster rate, yet R1 stabilized at younger ages. These developmental patterns reveal that myelin matures heterochronously in the frontal cortical ribbon, with early maturation in deeper cortical layers and protracted myelination in superficial layers.

### Frontal cortex myelin development reflects hierarchy and anatomy

In addition to uncovering divergent R1 maturation between superficial and deep cortical compartments, cortical maps of R1 change (**Fig. 3b, c**) showed considerable regional heterogeneity in R1 development within each cortical compartment. Quantifying the coefficient of variation (CV; dispersion around the mean) in the rate of R1 increase across frontal regions within each compartment revealed that deeper cortex showed relatively more uniform increases in R1 across regions, whereas middle and superficial cortex exhibited heightened regional variability in R1 development (superficial: CV = 0.48; middle: CV = 0.40; deep: CV = 0.31). We aimed to identify key contributors to regional developmental variability within each compartment, hypothesizing that asynchronous myelination may be explained by cortical hierarchical position^5,12–14,56^ and local cytoarchitecture^11^. To index each region’s hierarchical position in the frontal lobe, we used the hierarchical sensorimotor-association (S-A) axis of cortical organization^3^ (**Fig. 4a**). To capture regional differences in local anatomy, we used a main axis of cytoarchitectural variation derived using the BigBrain 3D histological atlas of cell body staining (**Fig 4d**).

**Figure 4.**
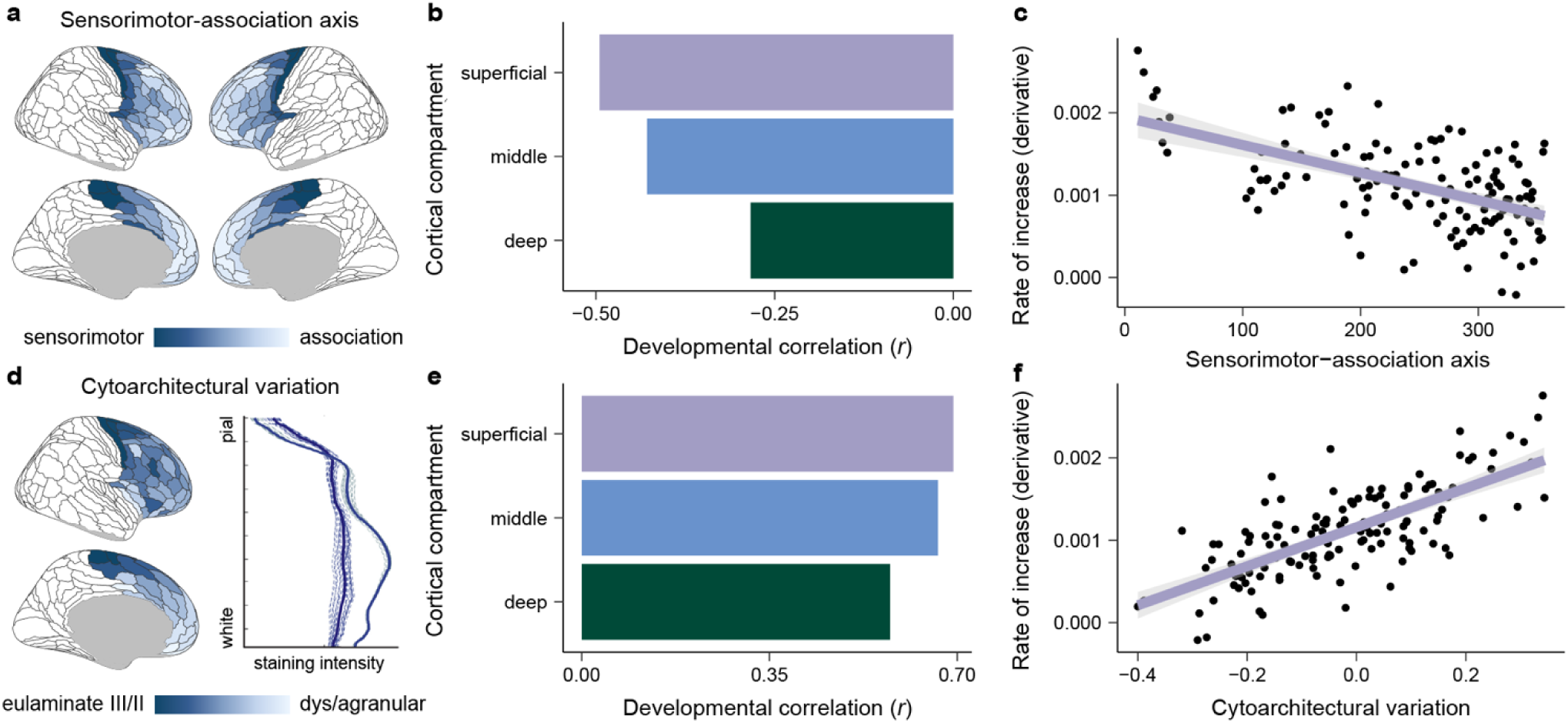
Superficial and deep R1 development is influenced by cortical hierarchy and cytoarchitectural anatomy. **a**) Each frontal lobe region is ranked along the sensorimotor-association (S-A) axis, a hierarchical axis of cortical feature derived in Sydnor et al.^3^. **b**) Developmental correlations quantifying the association between regional rates of R1 increase (average first derivative) and regional S-A axis ranks within the frontal lobe are presented for superficial, middle, and deep cortical compartments. Correlations were negative, indicating that the average rate of R1 increase was largest at the motor pole of the frontal S-A axis and progressively declined towards its association pole. The strength of the negative correlation increases from deep to superficial cortex. **c**) A scatterplot of the developmental correlation with the S-A axis in the superficial cortical compartment illustrates hierarchical myelination of superficial cortices during youth. **d**) Each frontal lobe region is ranked along an axis of cytoarchitectural variation obtained by Paquola et al. ^57^ using the BigBrain 3D histological atlas, a post-mortem atlas of cell body staining. Cytoarchitectural variation was characterized by cross correlating depth-wise straining intensity profiles between different cortical locations, thereby indexing regional differences in neuron density and soma size throughout the cortical ribbon. Cytoarchitectural variation captures a continuum of cortical types along a eulaminate to dys/agranular axis characterized by decreasing laminar differentiation^15^. **e**) Developmental correlations quantifying the association between regional rates of R1 increase and position on the frontal axis of cytoarchitecture variation are shown for superficial, middle, and deep cortical compartments. Correlations were robust and positive across compartments, linking accelerated myelination to eulaminate cortices and slower myelination to dys/agranular cortices. **f**) A scatterplot of the developmental correlation with the cytoarchitectural axis is displayed for the superficial cortical compartment, highlighting differences in R1 development depending on a region’s anatomical profile.

We observed negative correlations between regional rates of R1 increase and regional S-A axis ranks in superficial, middle, and deep cortex, signifying that the rate of myelination was always greatest in primary and early motor cortex and progressively decreased across regions positioned higher in the frontal cortical hierarchy. However, the magnitude of developmental correlation with the S-A axis was considerably greater in superficial and middle cortical compartments than in deep cortex (superficial: *r* = -0.49; middle: *r* = -0.43; deep: *r* = -0.28; **Fig. 4b, c**). In contrast, correlations between regional rates of R1 increase and the axis of cytoarchitectural variation were quite strong in all compartments (superficial: *r* = 0.69; middle: *r* = 0.66; deep: *r* = 0.57; **Fig. 4e, f**), linking regional differences in R1 change to microstructural anatomy throughout the cortical ribbon.

These compartment-wise correlations imply that contributors to asynchronous regional myelin development may differ in deep and superficial cortex. To directly test this idea, we fit multiple regressions with the S-A axis and cytoarchitectural axis as joint predictors of regional rates of R1 change in superficial, middle, and deep cortex. In superficial and middle cortex, both the S-A axis (superficial: *t*-*value* = -3.12, *p* = 0.002; middle: *t*-*value* = -2.53, *p* = 0.013) and the cytoarchitectural axis (superficial: *t*-*value* = 8.63, *p* < 0.001; middle: *t*-*value* = 8.31, *p* < 0.001) were significantly associated with regional rates of R1 increase. In deep cortex, however, the cytoarchitectural axis was significantly associated with regional R1 development (*t-value* = 6.97, *p* < 0.001), yet the S-A axis was not (*t-value* = -0.58, *p* = 0.561). Cortical features linked to the heterogeneous growth of a plasticity regulator thus differ laminarly. Asynchronous myelination between regions with distinct cytoarchitectures is present throughout the cortical ribbon, whereas hierarchically patterned myelination is characteristic of superficial layers that express heightened developmental variability.

### Laminar myelin trajectories differ between functionally diverse regions

Thus far, we have identified two key processes governing myelin development in the frontal cortex: myelin matures heterochronously across superficial and deep cortical compartments as well as heterogeneously within each cortical compartment. We next endeavored to characterize how these simultaneously evolving processes interact to govern laminar trajectories of myelin maturation in different areas of the frontal lobe. To enable this characterization, we ran a principal component analysis (PCA) to elucidate the spatial topography of regional differences in the shape of compartment-wise R1 developmental trajectories. The input to this PCA was each region’s GAMM-derived age splines from superficial, middle, and deep cortical compartments. The first principal component output by this PCA represents the principal spatial component of maturational variability in laminar R1 development (**Fig. 5a**).

**Figure 5.**
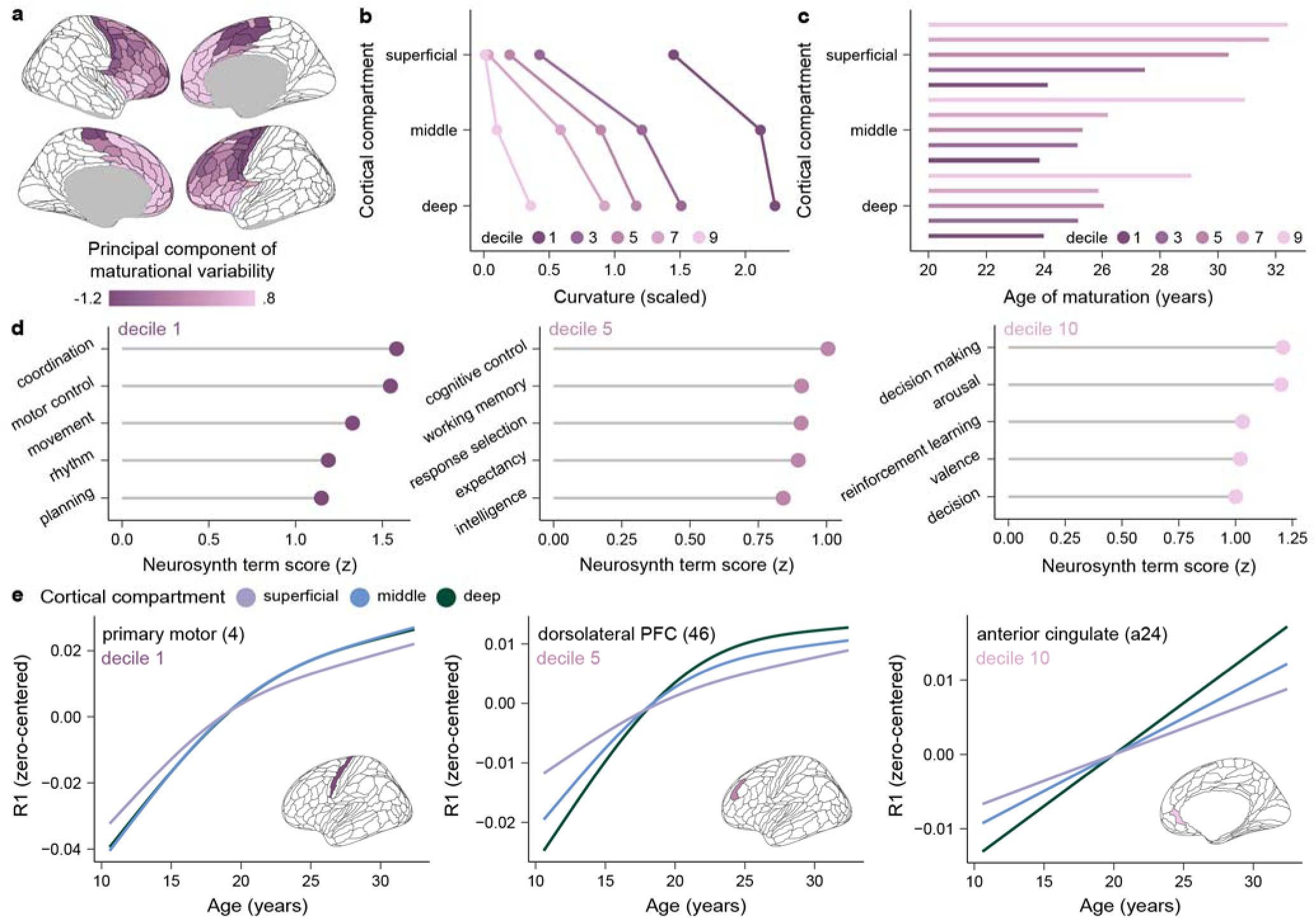
Heterochronous R1 maturation is differentially expressed in functionally distinct regions. **a**) Principal component analysis applied to compartment-wise developmental trajectories identified the principal component of maturational variability in laminar R1 development. **b**) Frontal regions located in different deciles of the principal component of maturational variability exhibit R1 age splines with varying patterns of curvature across superficial, middle, and deep cortical compartments. High curvature values index non-linear, curved developmental splines with clear plateaus. Low curvatures values signify relatively more linear spline fits. Curvature was calculated based on the shape of GAMM-derived age splines. **c**) Frontal regions located in different deciles of the principal component of maturational variability exhibit differing degrees of maturational heterochronicity across superficial, middle, and deep cortex. **d**) Neurosynth-based functional decoding identifies the primary functions supported by frontal regions included in the first, fifth, and tenth decile of the principal component of maturational variability. Functional decoding was performed using meta-analytic maps of over 100 psychological terms derived from prior task-based functional MRI studies. **e**) Developmental trajectories of R1 (zero-centered GAMM splines) are shown in superficial, middle, and deep cortex for exemplar frontal regions falling within the first, fifth, and tenth decile of the principal component of maturational variability. Regional plots illustrate different modes of laminar R1 development.

The first principal component explained 88% of the variance in R1 developmental profiles, capturing a cortical spectrum of laminar myelin maturation (**Supplementary Fig. 5.1**). Frontal regions located in different positions along this spectrum had R1 age splines with different curvature patterns (**Fig. 5b**) that plateaued at different maturational ages (**Fig. 5c**) across superficial, middle, and deep cortex. To further unpack these differences, we divided the principal component of maturational variability into deciles and probed decile-specific functional and developmental properties. The first decile (dark pink in **Fig. 5)** comprised regions necessary for motor control and coordination (**Fig. 5d**, left). Within this motor-related decile, superficial, middle, and deep cortex exhibited R1 trajectories with high curvatures values that matured simultaneously in the early 20s, as seen in the primary motor cortex (**Fig. 5e**, left). The fifth decile (medium pink in **Fig. 5**) was composed of lateral PFC regions that support cognitive control and working memory (**Fig. 5d**, middle). In this cognition-linked decile—well exemplified by the dorsolateral PFC (**Fig. 5e**, middle)—R1 age splines had high curvature and matured in the mid-20s in middle and deep cortical compartments, yet exhibited protracted increases into the 30s in superficial cortex. Finally, the tenth decile (light pink in **Fig. 5**) was functionally linked to decision making, arousal, and valence and localized to the anterior insula, medial PFC, and anterior cingulate (**Fig. 5d, e**, right). Frontal regions in this decile had low-curvature age splines that showed continuous increases in R1 into the 30s across superficial, middle, and deep cortex. These results underscore how divergent maturational timing between superficial and deep layers is differentially expressed in functionally diverse areas of the frontal cortex, with heterochronicity being most pronounced in lateral PFC regions.

### Developmental results are robust in sensitivity analyses

To ensure that the observed patterns of frontal cortex R1 development applied across sexes and were robust to potential biological or methodological confounds, we fit sex-stratified models and additionally performed a series of sensitivity analyses. R1 did not significantly differ between females and males in any region in superficial, middle, or deep cortex (*p*_FDR_ > 0.05 for the main effect of sex in all models). R1 developmental splines also did not significantly differ by sex (*p*_FDR_ > 0.05 for all age-by-sex interactions), indicative of similar developmental trajectories in females and males. Moreover, when covarying developmental models for biological sex, developmental patterns were unchanged compared to main analyses.

We subsequently evaluated whether age-dependent change in R1 within the frontal cortical ribbon replicated when covarying compartment-specific GAMMs in each region for cortical thickness, cortical gyrification (indexed by mean curvature), structural data quality (indexed by the euler number), and cortical partial voluming (indexed by the cortex volume fraction). In all sensitivity analyses, R1 significantly increased in the vast majority of frontal regions (>88%) in all three cortical compartments. Moreover, in all of the sensitivity analyses performed, both compartment-wise and regional patterns of the rate of R1 increase and the age of R1 maturation very closely mirrored results from the main analysis (**Fig. 6a, b**). Finally, regardless of the biological or methodological covariate included in sensitivity analyses, different modes of laminar R1 maturation could be seen within functionally diverse cortical regions (**Fig. 6c**) and the principal component of maturational variability was highly stable (**Supplementary Fig. 6.1**). Accordingly, our findings concerning R1 development within the frontal cortical ribbon are consistent across sexes and are not driven by individual differences in cortical architecture or structural data quality.

**Figure 6.**
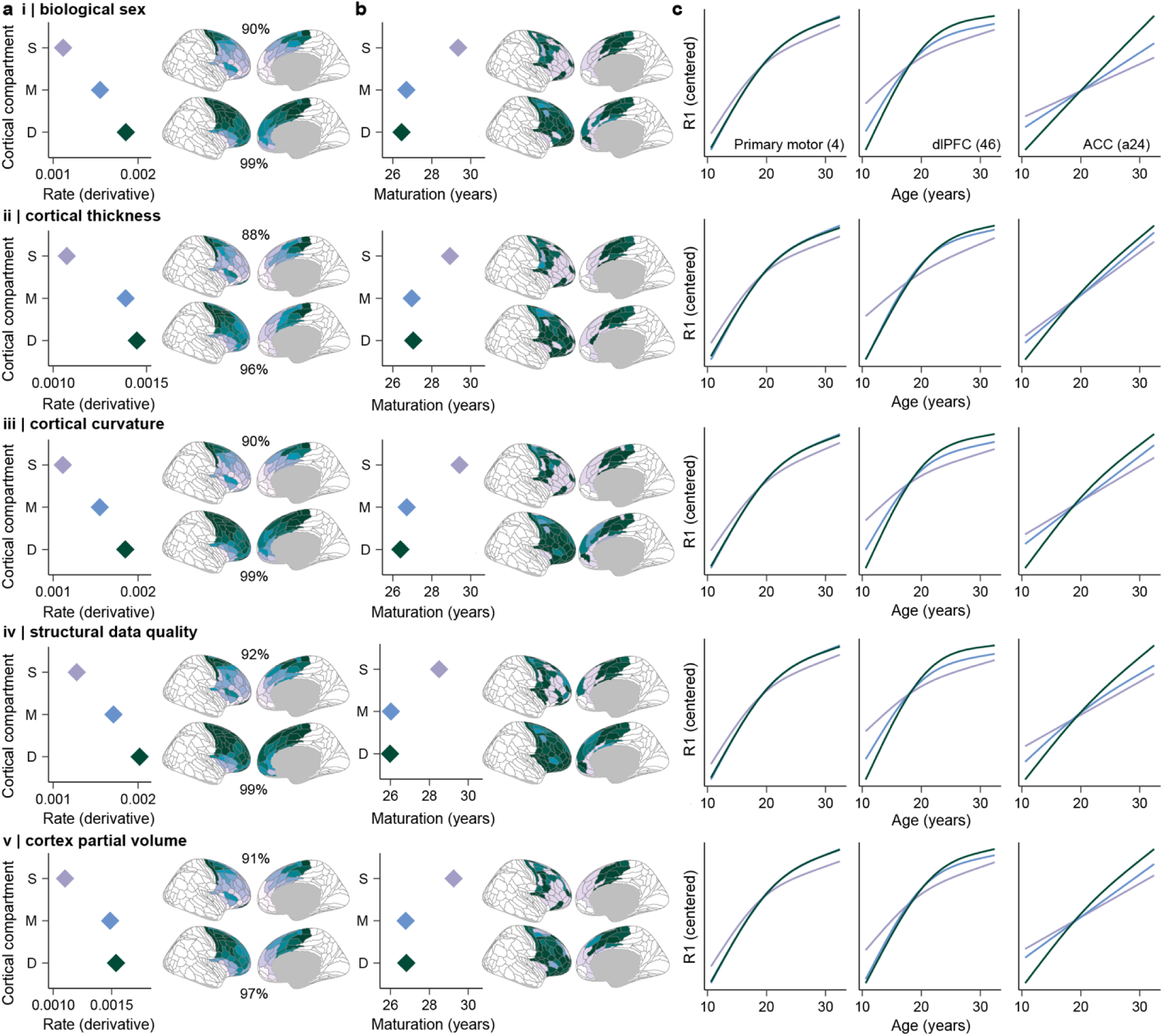
R1 developmental patterns are robust in sensitivity analyses. Regional and compartment-wise patterns of frontal cortex R1 development are robust to controls for individual differences in i) biological sex, ii) cortical thickness, iii) cortical curvature, iv) structural data quality, and v) cortex partial volume effects. Key results are shown for each of these five sensitivity analyses; all results strongly converge with findings from the main analysis. **a**) Compartment plots display the average rate of R1 increase in the frontal lobe in superficial, middle, and deep cortex. The corresponding cortical maps present the average rate of R1 increase in each frontal region in superficial cortex (top) and deep cortex (bottom). Percentages indicate the percent of frontal lobe regions that exhibited a significant increase in R1 in the corresponding compartment. **b**) Compartment plots chart the average age at which R1 matured in the frontal lobe in superficial, middle, and deep cortex. The corresponding cortical maps show the age of R1 maturation in each frontal region in superficial cortex (top) and deep cortex (bottom). **c**) Developmental trajectories of R1 are shown in superficial, middle, and deep cortical compartments for the primary motor cortex (area 4), the dorsolateral prefrontal cortex (dlPFC; area 46) and the anterior cingulate cortex (ACC; area a24).

### Frontal cortex myelin refines E/I-linked neural dynamics

Continued myelination of the developing frontal cortex over the span of decades should have a profound, yet under-characterized, effect on cortical circuit physiology. Given that the majority of cortical myelin forms around afferent and efferent excitatory connections and inhibitory PV interneurons^24–26,58^, we theorized that intracortical myelin would alter the local E/I ratio and E/I-linked neural dynamics. To test this theory, we related intracortical R1 to aperiodic EEG activity, which provides a readout of non-oscillatory, population-level neural activity. Computational, pharmacological, and neurochemical data have shown that decreases in the aperiodic exponent—a measure of how power decays with increasing frequency—reflect increases in E/I balance^59,60^ and a faster timescale of neural activity^61,62^. We therefore predicted that greater R1 in the cortex would be linked to a lower aperiodic exponent, providing evidence that myelin balances fast pyramidal^36^ and PV neuron firing^36,49^ to facilitate faster dynamical changes in population activity.

We source-localized 64-channel scalp EEG to each individual’s cortical surface and calculated the aperiodic exponent within individual frontal lobe regions (**Fig. 7a-c**). Replicating prior findings^60,63,64^, the aperiodic exponent significantly declined with age across the frontal cortex (indicative of a flattening of the aperiodic slope; *F* = 16.7; *p* < 0.001; **Fig. 7d**). Given that pyramidal neurons in both superficial and deep cortical layers substantially contribute to scalp-recorded EEG^65,66^, we aimed to study not only whether R1 was related to the aperiodic exponent, but also whether the strength of this relationship differed for R1 measured in superficial and deep cortex. We investigated these aims simultaneously by fitting region-specific GAMMs with both main and interaction effects (controlling for age): the main effect characterized the association between the exponent and deep cortex R1 while the interaction term tested whether the association significantly differed in strength in superficial cortex.

**Figure 7.**
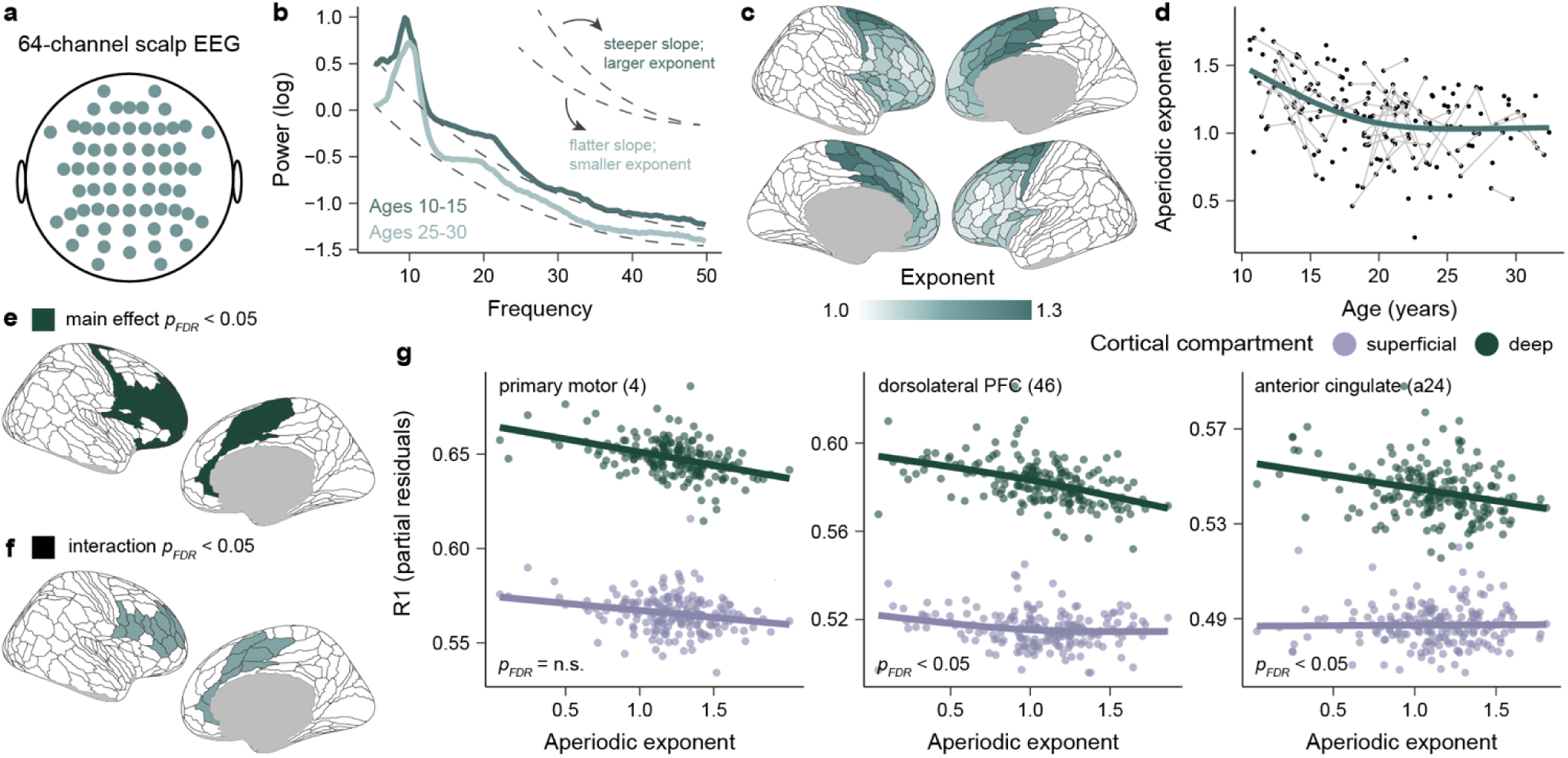
Frontal cortex R1 is associated with EEG neural dynamics. **a**) Neural activity was measured across the cortex using 64-channel scalp EEG. **b**) The aperiodic exponent was calculated from the slope of the 1/f power spectral density (approximated by the dotted line for illustration). The exponent quantifies how quickly power decays with increasing frequency. The average power spectrum across frontal lobe EEG electrodes is shown for individuals 10-15 and 25-30 years old. **c**) The EEG aperiodic exponent is shown source localized to individual frontal lobe regions. Source reconstruction was used to map each participant’s preprocessed EEG activity to their individual-specific cortical surface prior to parameterizing aperiodic activity in each frontal region. The exponent was averaged across participants in each region for visualization. **d**) The frontal cortex aperiodic exponent (averaged across all frontal regions) decreases during adolescent neurodevelopment. The teal line represents the GAMM-derived developmental trajectory overlaid on participant-level data with longitudinal EEG sessions connected. **e**) A cortical map displaying frontal regions that exhibited a significant association between deep cortex R1 and the aperiodic exponent. **f**) A cortical map highlighting frontal regions where the strength of the R1-exponent relationship significantly differed in deep and superficial cortex. **g)** Interaction models uncovered significantly stronger negative associations between the aperiodic exponent and R1 measured in deep as compared to superficial cortex. The relationship between the aperiodic exponent and R1 within both superficial and deep cortex is shown for the primary motor cortex, the dorsolateral prefrontal cortex (PFC), and the anterior cingulate. R1 values are partial residuals adjusted for age. *p*_FDR_ values represent the significance of the exponent-by-compartment interaction.

The aperiodic exponent was significantly (*p*_FDR_ < 0.05) related to deep cortex R1 in regions spanning both lateral and medial portions of the frontal lobe (**Fig. 7e**). As predicted, higher R1 was associated with a lower exponent. In the majority of regions with a significant main effect, the interaction term was also significant (*p*_FDR_ < 0.05; **Fig. 7f**), indicating that the nature of the R1-exponent relationship differed between superficial and deep cortical compartments. Probing these interactions revealed that the strength of the R1-exponent relationship was significantly stronger for R1 measured in deep than in superficial cortex (**Fig. 7g**). Altogether, the present results link greater myelin content, particularly within deep cortical layers, to an EEG signature of faster cortical dynamics.

### Frontal cortex myelin supports faster learning and processing speed

In a final set of analyses, we explored whether myelin in superficial computation layers and deep output layers supports unique aspects of efficient cognitive processing. We predicted that myelination of superficial, cortico-cortical, computational circuits would facilitate rapid learning^36,37^, whereas myelination of deep, subcortically-projecting, output circuits would support faster output processing speed. To test these predictions, we derived measures of learning rate and processing speed from a youth-friendly sequential decision-making task^67,68^ that involved learning stimulus-outcome relationships to earn rewards in two task stages. Stages were characterized by fixed (stage 1) and dynamically changing (stage 2) stimulus-outcome probabilities (**Fig. 8a**). To operationalize learning, we applied a Bayesian reinforcement learning model to trial-level data and calculated participant learning rates on each stage of the task. To index processing speed, we calculated the average response time on each stage of the task. Both learning rates (*paired t* = 2.43, *p* = 0.016) and response times (*paired t* = 14.76, *p* < 0.001) significantly increased within-person between stages 1 and 2 of the task (**Fig. 8b**), signifying that stage 2 elicited a faster speed of updating and slower decision making due to its changing (as opposed to stable) reward contingencies. We studied how task performance changed throughout development using both group-level GAMMs and measures of within-individual change. In group-level GAMMs, learning rates slightly increased with age in a linear fashion (stage 1: *F* = 5.51, *p* = 0.020; stage 2: *F* = 8.81, *p* = 0.004; **Fig. 8c**) and response times substantially decreased with age during the adolescent period (stage 1: *F* = 29.98, *p* < 0.001; stage 2: *F* = 7.85, *p* = 0.005; **Fig. 8d**). In a complementary analysis that used person-mean centered regression to test for within-individual longitudinal change, response times significantly decreased within-person with age (stage 1: *t-value* = -2.13, *p* = 0.037; stage 2: *t-value* = -3.90, *p* < 0.001); within-person changes in learning rates were not significant.

**Figure 8.**
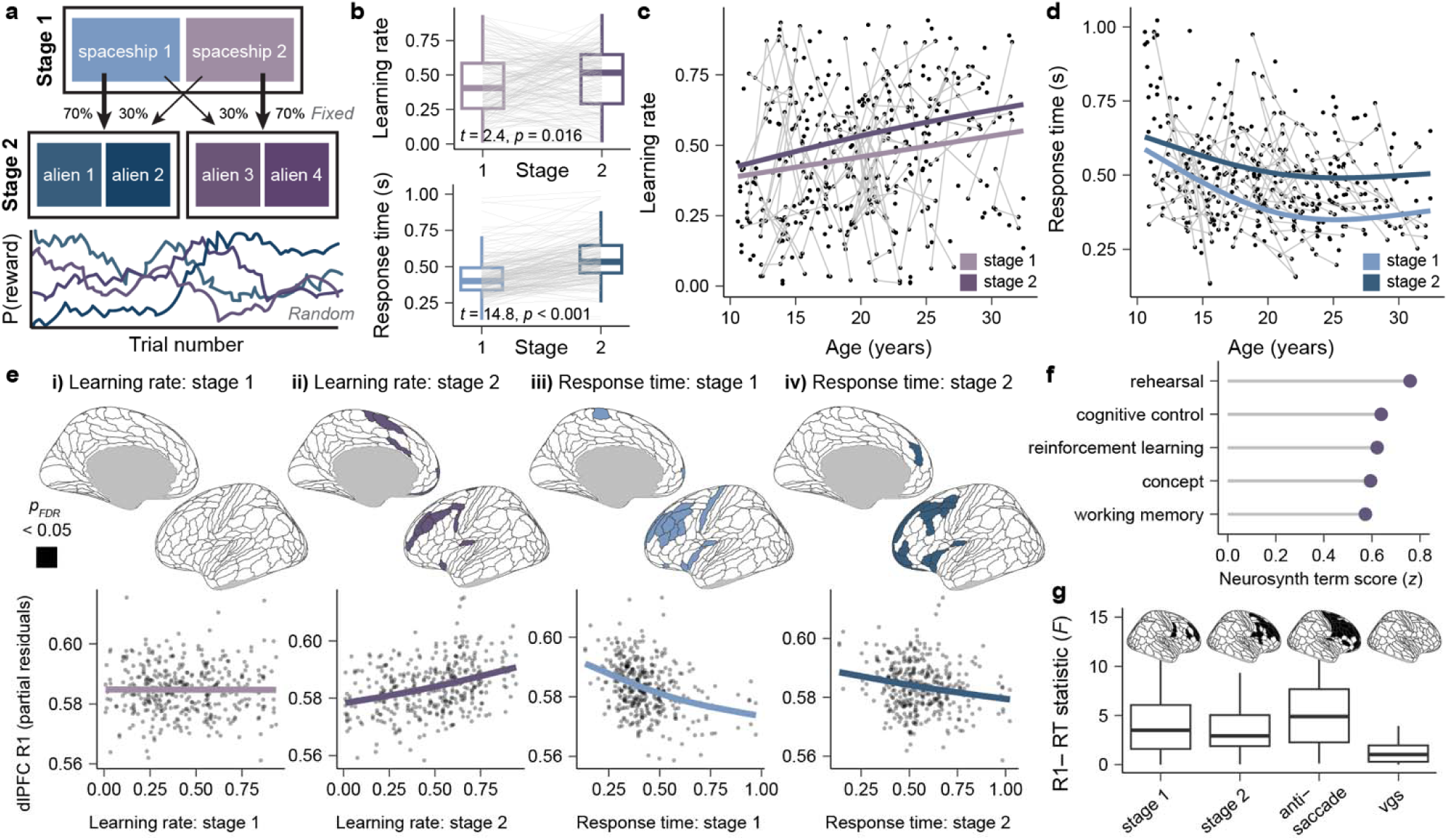
Prefrontal cortex R1 is linked to learning rate and processing speed. **a**) Participants completed a two-stage sequential decision-making task to earn rewards. During the first stage of each trial, participants selected one of two spaceships that transitioned with fixed probabilities to either a common planet (70% of transitions) or rare planet (30% of transitions). During the second stage of each trial, participants selected one of two aliens on the planet for the chance to receive a reward; the probability of receiving a reward (P(reward)) from each alien fluctuated randomly to incentivize continuous learning of changing contingencies. **b**) Learning rates and response times significantly increased within-person (grey lines) between stages 1 and 2 of the task, as assessed with paired-samples t-tests. **c**) Learning rates were variable across individuals of all ages during both task stages. **d**) Response times decreased non-linearly with age on stage 1 and 2 of the task. **e**) Cortical plots are shown that highlight frontal regions where R1 was significantly associated with i) stage 1 learning rates, ii) stage 2 learning rates, iii) stage 1 response times, and iv) stage 2 responses times. Significant regions are colored. Below each cortical plot, the relationship between each cognitive measure and R1 in the dorsolateral prefrontal cortex (dlPFC) is shown in a partial residual plot. **f**) Neurosynth-based functional decoding of brain regions that exhibited a significant association between R1 and stage 2 learning rates (purple regions in eii) emphasizes their role in cognitive control and reinforcement learning processes. **g**) Associations between frontal R1 and response times (RT) were larger when engaging higher-order cognitive control (stage 1, stage 2, anti-saccade) than when performing a simple visual-motor response (visually-guided saccade, vgs). The cortical plots highlight frontal regions with a significant relationship between R1 and task-specific response times; significant regions are in black. Boxplots summarize the distribution of regional *F*-statistics derived from GAMMs relating frontal R1 to RT on each task.

As in the analyses linking R1 to EEG aperiodic activity, we used GAMMs to simultaneously study compartment-specific associations between R1 and cognitive measures (main effects) and whether the strength of R1-cognition associations differed between R1 in superficial and deep cortical compartments (interaction effects). However, interaction terms were not significant in any frontal region for any cognitive measure (*p*_FDR_ > 0.05 for all interactions tested), suggesting that R1 in superficial and deep cortex was similarly related to task performance. We therefore combined across superficial and deep cortical compartments for all further analyses, deriving a single relationship between task measures and R1 in the cortical ribbon. Examining associations between regional R1 and learning rates revealed that no frontal region showed a significant relationship between R1 and learning rate during stage 1 of the task (**Fig. 8ei**). In contrast, during stage 2 of the task, faster learning rates were significantly associated with higher R1 in 16 frontal regions (*p*_FDR_ < 0.05). Significant regions primarily localized to the dorsolateral PFC (**Fig. 8eii**) and comprised cortices recruited by tasks that involve cognitive control, working memory, and reinforcement learning (**Fig. 8f**). Greater R1 in higher-order cognitive regions was thus linked to faster learning during task stages characterized by rapidly evolving—but not stable—reward contingencies.

Models relating R1 to response times showed that greater R1 in the lateral PFC was significantly associated with faster responding during both stage 1 of the task (29 significant regions at *p*_FDR_ < 0.05; **Fig. 8eiii**) and stage 2 of the task (37 significant regions at *p*_FDR_ < 0.05; **Fig. 8eiv**). We theorized that these broadly distributed relationships may reflect the capacity for prefrontal myelin to enhance cognitive processing speed, rather than just the speed of motor outputs. To test this theory, we used GAMMs to relate R1 to response times on an anti-saccade task—a prefrontal-dependent task that taxes inhibitory control—and a visually-guided saccade task—a task that gauges sensorimotor processing speed. This allowed us to compare results from the decision-making task to independent tasks of intentional cognitive control and reflexive visual-motor responding. Faster responding on correct trials of the anti-saccade task was associated with significantly higher R1 across much of the frontal lobe (71 significant regions at *p*_FDR_ < 0.05). Faster responding on the visually-guided saccade task was not, however, significantly associated with R1 in any frontal region. These specificity analyses (**Fig. 8g**) demonstrate that greater myelination of the prefrontal cortex supports faster processing speed specifically when endogenously engaging higher-order cognitive processes.

## Discussion

During the process of embryonic corticogenesis, layers of the cerebral cortex form in an inside-out manner; *ex vivo* studies have shown that layers 2/3 experience later windows of neurogenesis and laminar differentiation than layers 5/6^69^. In the current study, we demonstrate *in vivo* that the maturation of myelin follows a similar sequence during youth. In doing so, we extend a fundamental principle of prenatal cortical formation to postnatal cortical plasticity. We show that myelin-sensitive R1 matures heterochronously between superficial and deep compartments of the prefrontal cortical ribbon, with relatively more superficial cortex exhibiting shallower and more protracted increases in R1 than deep cortex. This cortical depth-related divergence in maturational timing is differentially expressed across the frontal lobe, resulting in different profiles of laminar myelin development between cytoarchitecturally and functionally distinct regions. By combining myelin-sensitive MRI with measures of EEG aperiodic activity, we uncovered that higher R1 was related to a functional index of higher E/I balance^59,60^ and a faster timescale of neural activity^61,62^. Finally, using tasks of decision making, inhibitory control, and oculomotor saccades, we linked higher R1 in both deep and superficial compartments of the lateral PFC to enhanced learning rates and faster cognitive—but not sensorimotor—processing speed. Together, these results provide evidence for asynchronous reductions in plasticity in deep “output” layers and superficial “computation” layers of the human PFC. This asynchronous maturation likely allows the developing association cortex to couple cognitively-relevant increases in circuit stability and efficiency with extended neuroplasticity.

The diverging trajectories of R1 development observed in superficial and deep compartments of the prefrontal cortical ribbon provide insight into laminar variation in the timing of myelin maturation and the patterning of developmental plasticity. In deeper cortex proximal to layers 5/6, increases in R1 were larger, plateaued earlier, covaried with differences in cytoarchitecture, and occurred relatively more synchronously across frontal regions. In superficial cortices that localize approximately to layers 2/3, increases in R1 were shallower, protracted, and expressed with heightened and hierarchically-organized topographic variability. Together, these data provide evidence for earlier maturation of layer 5/6 cortical-subcortical and feed-back cortical projections, coupled with extended plasticity of layer 2/3’s highly integrative cortico-cortical circuitry^19,70^. Preferential myelination of deep PFC layers may function to consolidate top-down connectivity earlier during youth, ensuring that pyramidal cells that produce behavioral outputs propagate signals reliably and quickly. In contrast, shallow yet prolonged myelination of superficial PFC layers likely allows high-dimensional computational circuits to retain enhanced flexibility important for hierarchical refinements in learning and memory^36,71^. Notably, this laminar divergence in R1 development was differentially expressed across frontal regions. Laminar heterochronicity was absent in frontal motor regions, evolved over the longest period of time in medial prefrontal, insular, and anterior cingulate cortices, and was most pronounced in evolutionarily-expanded cognitive control regions in the lateral PFC.

Heterochronous myelination across superficial and deep layers of the PFC may reflect differences in the degree to which myelin growth results from activity-independent genetic programs versus experience-dependent increases in neuronal activity^33,58,71–73^. Although oligodendrocyte precursor cells are evenly dispersed throughout cortical layers, myelinating oligodendrocytes increase in density in layers 5/6^58,72,74^. This cell type distribution results from an intrinsic capacity for pyramidal neurons in deep layers to stimulate oligodendrocyte differentiation and myelin biogenesis independent of activity^58^. In contrast, pyramidal neurons in superficial layers 2/3 engage in highly recurrent excitatory dynamics^19,20^ sufficient to trigger modes of myelin formation that are coupled to neuronal activity. Moreover, superficial layers wrap a greater proportion of total myelin around PV interneurons, which undergo extensive activity-dependent myelination in response to environmental inputs during youth^75,76^. The shift from earlier-maturing, synchronous R1 increases in deep cortex to protracted, hierarchical R1 increases in superficial cortex may thus reflect a transition from genetically-programmed to experience-driven myelin formation. This shift would allow the cortex to both establish a reliable foundation of myelin necessary for rapid cortico-subcortical signaling and to continuously myelinate cortical connections in response to evolving environmental demands.

Although prolonged myelination of the cortex is a hallmark of postnatal development, how it impacts the function of local cortical circuits remains an active area of exploration. We found that greater R1—especially within deeper cortical layers—was associated with a smaller exponent of the aperiodic component of EEG, linking higher myelin content to more mature patterns of neuronal activity^60,64^. The aperiodic exponent decreases with a desynchronization of population-level firing^77^ that results in faster temporal fluctuations in (i.e., a faster timescale of) activity^61,62^. The aperiodic exponent is also strongly modulated by local E/I interactions^59,60^, together indicating that myelination of pyramidal and PV cells may alter E/I balance and activity dynamics^49^ to allow maturing circuits to signal with higher temporal efficiency. Laminar electrode recordings have shown that the aperiodic exponent decreases from superficial to deep cortex, reflecting a faster timescale of activity in output layers 5/6 than in computational layers 2/3^62^. The present results linking R1 to the aperiodic exponent suggest that pronounced developmental increases in myelin in deep cortex may help establish the fast temporal dynamics observed in cortical output layers^20,78^. In contrast, gradual myelination of superficial cortex may result in an uncoupling of its function from local anatomy and produce the sustained, recurrent dynamics characteristic of computational circuitry^19,20^.

Rodent studies have demonstrated that activity-dependent myelin sheath formation occurs in the cortex on learning-activated axons^37,38^, supporting the hypothesis that cortical myelin content may impact learning capacity in humans. Consistent with this hypothesis, we found that higher R1 within the dorsolateral PFC was associated with a higher learning rate, particularly when the task demanded continued updating and behavioral flexibility. In complementary experiments, higher R1 in the PFC was associated with faster responding when engaging higher-order cognitive control but not lower-order visuomotor processes. Together, these findings signify that myelination of the human PFC may support age-related improvements in learning and cognitive processing speed, thereby enhancing overall cognitive efficiency. Contrary to our prediction, R1 associations with learning rate were not confined to superficial cortex, nor were processing speed associations specific to deep cortex. This lack of laminar specificity may reflect the fact that both behavioral readouts required the coordinated engagement of superficial computational and deep output circuitry. In particular, our computational measure of learning required producing motor outputs and our measure of processing speed depended on computing a decision. Future studies may clarify whether myelin in layers 2/3 and 5/6 uniquely contributes to distinct behavioral operations by tracking longitudinal change in superficial and deep myelin following intensive learning or motor training.

This study incorporates methodological innovation through the use of myelin-sensitive imaging, intracortical depth profiling, and 7T longitudinal MRI; however, there are several methodological limitations. First, although myelin is the main source of R1 signal in the cortex, iron makes a non-negligible contribution to cortical R1^79^. Importantly, however, most cortical iron co-localizes with oligodendrocytes and myelinated intracortical axons, thus laminar variation in iron provides indirect insight into the relative density of myelin^80,81^. Second, the spatial resolution of the R1 data used here is not sub-millimeter, although it is equivalent to the resolution used in prior studies that sampled myelin-sensitive imaging measures at 10 or more cortical depths^11,41^. This resolution smooths sharp changes in laminar myelin profiles (e.g., turning points at layer boundaries)^82^ and can result in the smoothing or blurring of R1 signal across spatially proximal layers and compartments.

Third, equidistant depth sampling can be used to differentiate between superficial and deep cortical compartments that spatially approximate layers 2/3 and 5/6, but the mapping between compartments and layers is inexact and may vary across regions with distinct laminar architectures. Fourth, due to the instrumental use of ultra-high field and quantitative MRI, the size of the study sample is considerably smaller than many publicly available imaging samples collected at 3T, although the sample is one of the largest pediatric 7T samples collected to date.

During childhood and adolescence, the maturation of the human cortex is governed by large-scale spatial axes that organize the temporal unfolding of plasticity across cortical regions^3,8,9,12,23,28,84^. The present study suggests that differences in temporal windows of plasticity also emerge between superficial and deep layers within the developing PFC. Divergent maturational timing across superficial and deep layers is an underrecognized mechanism through which late-developing association cortices balance increases in circuit robustness with protracted adaptability. Critically, extended malleability in superficial layers may increase their susceptibility to developmental and environmental insults—contributing to an outsize role of layer 2/3 circuits in transdiagnostic psychopathology^53,85,86^. Moving forward, continued elucidation of regional and layer-specific time windows of plasticity will allow for the precise demarcation of circuit-specific critical periods of vulnerability and adaptability.

## Methods

### Developmental dataset

This study analyzes data from a healthy, accelerated longitudinal youth sample recruited from the greater Pittsburgh area by the Laboratory of Neurocognitive Development at the University of Pittsburgh. Individuals in this sample participated in one to three longitudinal sessions scheduled approximately 20 months apart. Each session consisted of three visits, including a cognitive testing visit, a 7T MRI visit, and an EEG visit, which typically occurred on different days over the course of a few weeks. Study exclusion criterion included contraindications to receiving an MRI, an IQ below 80, a history of loss of consciousness due to head injury, a history of substance abuse, and a history of major psychiatric or neurological conditions for the participant or a first-degree relative. All individuals over the age of 18 years gave written informed consent before study participation. Individuals under the age of 18 gave informed assent along with written parental consent. Participants received monetary compensation for participation in the study. All study procedures were approved by the Institutional Review Board at the University of Pittsburgh.

Following data quality exclusions (see Sample construction below), data from 140 individuals and 215 longitudinal imaging sessions were included in this study (85 individuals with 1 session, 35 individuals with 2 sessions, 20 individuals with 3 sessions). The inter-scan time interval between imaging sessions 1 and 2 was 21.6 months (± 4.2 months) on average, with a mode of 20.5 months. The inter-scan interval between imaging sessions 2 and 3 was 23.2 months (± 7.3 months) on average, with a mode of 20.7 months. Participant demographics included an age range of 10 to 32 years (mean age = 20.0 years ± 5.4) and a self-reported sex assigned at birth breakdown of 70 females and 70 males. Participants (18 years and older) or their parents identified the study participant’s race/ethnicity: 8.6% of participants identified as Asian, 15% as Black, 2.9% as Hispanic, 67.1% as White, and 5.7% as multiracial; data were missing for 1 participant (remaining 0.7%).

National percentile rankings of the area deprivation index for this study sample ranged from 3^rd^ percentile to 99^th^ percentile with a mean of 50.0 (± 25.6).

### MP2RAGE acquisition

All MRI data were acquired on a single 7 Tesla Siemens Magnetom (MR B17) with parallel transmission (pTx). A Magnetization Prepared 2 Rapid Acquisition Gradient Echoes (MP2RAGE) sequence was collected and used to generate both a T1-weighted uniform (UNI) image and a quantitative T1 map, which were collectively used for intracortical myelin mapping. MP2RAGE sequences combine two gradient-recalled echo (GRE) images acquired with short (INV1; a predominantly T1w image) and long (INV2; a predominantly PDw image) inversion times. The combination of these images results in UNI and quantitative T1 images that are virtually free of B_1_^-^, proton density, and T2* effects and that have minimal B_1_^+^ transmit field inhomogeneity^87^. MP2RAGE data were acquired with an in-plane GRAPPA acceleration factor of 4 and the following parameters: inversion times of 800 ms (INV1) and 2700 ms (INV2), repetition time of 6000 ms, echo time of 2.87 ms, flip angles of 4 degrees (INV1) and 5 degrees (INV2), and a voxel resolution of 1 mm isotropic. Denoised UNI images with suppressed background noise were generated from this protocol using a robust (i.e., regularized; lambda = 10) combination of the two inversion time images^88^. Given the presence of enhanced B_1_^+^ field inhomogeneity present at ultra-high field strengths, MP2RAGE data were collected with a parallel transmit system to increase B_1_ ^+^ homogeneity and with an optimized protocol previously shown to minimize the sequence’s B_1_^+^ sensitivity^89^.

### MP2RAGE myelin imaging in the cortical ribbon

MP2RAGE sequences generate inherently co-registered UNI and quantitative T1 images. The UNI image can be used as a T1w image for segmentation and cortical surface reconstruction given its high gray-white contrast and contrast-to-noise ratio (CNR)^90^. The T1 map provides a direct quantitative readout of the longitudinal relaxation time in seconds. The longitudinal relaxation time represents the time it takes for the longitudinal magnetization to return to equilibrium along the direction of the main magnetic field, typically ranging from approximately 1.4 to 2.2 seconds in gray matter^89^. Due to the fact that the largest influence on T1 relaxation is the presence of lipids, which are both greatly enriched in and primarily accounted for by myelin^91^, the T1 map can be used for R1 (1/T1; longitudinal relaxation rate) myelin imaging. The presence of myelin reliably shortens the T1 relaxation time and correspondingly increases the longitudinal relaxation rate; as a result, R1 is higher in areas with greater myelin content^54,92,93^. R1’s sensitivity to cortical myelin concentration has been validated in multiple studies using anatomical mapping and post-mortem histology^44–47,93,94^. Furthermore, cortical R1 has been shown to have high CNR and excellent within-individual scan-rescan reliability at 7T, making it a valuable measure for use in longitudinal studies^48^.

UNI images and quantitative T1 maps were processed with the goal of mapping R1 values to different locations in the cortical surface. First, to mitigate the potential effects of residual B_1_^+^ transmit field inhomogeneity on the acquired structural data, the UNICORT (unified segmentation based correction of R1 brain maps for transmit field inhomogeneities) algorithm^95^ was applied to both denoised UNI images and raw quantitative T1 maps. UNICORT uses a probabilistic framework for simultaneous image segmentation, registration, and bias correction and was applied to UNI and T1 data using a containerized version of a statistical parametric mapping (SPM12) BIDS app. After UNICORT correction, UNI images were processed with FreeSurfer’s longitudinal processing stream to generate individual-specific cortical surfaces. FreeSurfer with run using a containerized FreeSurfer BIDS app (FreeSurfer version 7.4.1). Longitudinal FreeSurfer creates a within-participant template using robust, inverse consistent registration and initializes final structural processing using common information from the template^96^. This ensures that segmentation and surface reconstruction are performed consistently across, and unbiased to, individual timepoints, thereby enhancing longitudinal reliability^96^.

To enable cortical R1 mapping, volumetric R1 maps were computed from UNICORT-corrected quantitative T1 images (R1 = 1/T1 in sec^-1^) using 3dcalc from AFNI (version 23.1.10). R1 was then sampled to each participant’s timepoint-specific cortical surface using FreeSurfer’s volume-to-surface projection. R1 was sampled to surface vertices at 11 intracortical depths between the pial boundary and the gray-white boundary based on the relative fraction of cortical thickness. This allowed for the quantification of R1 at intracortical depths between 0% of cortical thickness (pial boundary) and 100% of cortical thickness (gray-white boundary) in 10% increments. Calculating depths based on percent of thickness ensures that R1 is sampled the same fraction into the cortex in all cortical regions, regardless of the region’s total cortical thickness. While a direct mapping between each cortical depth and specific cortical layers cannot be made, a general distinction between more superficial cortex (depths closer to the pial boundary) and deep cortex (depths closer to the white matter boundary) can be drawn. Furthermore, given that the relative proportion of the prefrontal cortical ribbon occupied by different layers has been described, approximate links can be drawn between depths <10% into cortex and layer 1, depths 10-45% into cortex and layers 2/3, and depths 65-100% into cortex and layers 5/6^19,43^, with the percent depth of layer 4 being notably variable across regions^97^.

After measuring R1 within different depths of cortex at all surface vertices, we quantified average R1 at each depth in individual cortical regions defined by the Human Connectome Project multi-modal parcellation (HCP-MMP)^98^. The HCP-MMP atlas was transformed from the fsaverage template to participants’ timepoint-specific cortical surfaces using FreeSurfer’s surface resampling algorithm, which employs spherical registration. Finally, we took analytic steps to ensure that we only analyzed R1 from cortical locations with 1) minimal to no partial voluming and 2) high quality R1 signal. To index partial voluming, we used participants’ FreeSurfer-processed UNI images and tissue class segmentations to calculate the cortex volume fraction for all 11 intracortical depths at every vertex. The cortex volume fraction quantifies the fraction of the signal estimated to come from cortical gray matter (as opposed to white matter, subcortical gray matter, or CSF). We only further analyzed 7 cortical depths where the average cortex volume fraction was >90%; these 7 depths ranged from 20% of cortical thickness to 80% of cortical thickness. To index signal quality, we computed the signal-to-noise ratio (SNR) of R1 in every HCP-MMP region, averaged across the 7 analyzed depths. We excluded 24 cortical regions in the temporal pole and medial prefrontal cortex with low SNR from all analyses. SNR was calculated as the vertex-wise mean of R1 divided by the standard deviation of R1 across all vertices included in a region.

### Sample construction

At the start of this study, identical parameter MP2RAGE scans (i.e., non-variant acquisitions as assessed with CuBIDS software^99^) were available for 264 imaging sessions collected from 159 participants. These data underwent additional curation to ensure that only highly quality neuroimaging data were analyzed. Data were excluded from this initially collected dataset for the following reasons: UNI images or R1 maps failed visual quality control for motion or scanner artifacts (44 scans excluded), imaging data were collected within 6 months of an included session (4 scans excluded), and the R1 map had a whole-cortex mean R1 > 4 standard deviations above the group-level mean of eligible scans (1 scan excluded). This sample construction procedure, which included both qualitative and quantitative quality assurance steps, led to a final dataset of 215 longitudinal sessions collected from 140 individuals.

### Brain maps of cortical anatomy, gene expression, and function

This study integrated previously published brain maps of cortical anatomy, gene expression, and function in order to explore convergence between MP2RAGE and independent myelin mapping techniques, contextualize developmental patterns, and perform functional decoding. The following cortical maps were used; each cortical map was parcellated with the HCP-MMP atlas using Connectome Workbench version 1.5.

#### T1w/T2w ratio map

The T1w/T2w ratio was measured throughout the cortex by Baum et al.^12^ in data collected from youth ages 8-21 years old as part of the Human Connectome Project in Development. T1w/T2w data were generated using HCP pipelines (PreFreeSurfer, FreeSurfer, and PostFreeSurfer pipelines) and corrected for B_1_^+^ bias using an empirically validated pseudo-transmit field correction approach^100^.

#### Magnetization transfer saturation

Magnetization transfer saturation measures the exchange of magnetic saturation from macromolecules—including those present in myelin—to protons in free water. Magnetization transfer saturation data were obtained from Hettwer et al.^41^, who derived this measure in youth ages 14-16 years old using a multi-parametric mapping sequence.

#### Myelin basic protein gene expression

A cortical map of myelin basic protein gene expression was produced by Wagstyl et al.^52^ by processing postmortem microarray data from the Allen Human Brain Atlas. Of note, Wagstyl et al. generated and validated dense, across-donor gene expression maps for the left cortical hemisphere only.

#### The sensorimotor-association axis

The S-A axis^3^ is a hierarchical axis of cortical organization derived from brain maps that provide complementary yet spatially correlated markers of cortical hierarchy. In particular, the S-A axis was derived by integrating *in vivo* markers of the cortex’s anatomical^101,102^, functional^103,104^, and evolutionary^105^ hierarchies as well as representative hierarchical gradients of spatial feature variation^106^. In the frontal lobe, the S-A axis spans from primary and early motor cortex to heteromodal and paralimbic association cortex.

#### A histology-based axis of cytoarchitectural variation

The axis of cytoarchitectural variation captures regional differences in depth-wise cytoarchitectural properties and spans from eulaminate cortex to dysgranular and agranular cortex. The cytoarchitectural axis was derived by Paquola et al.^57^ using depth-wise cell body (Merker) staining intensity profiles from the BigBrain 3D histological atlas. Diffusion map embedding was applied to a region-by-region correlation matrix of depth-wise staining intensity profiles (sampled at 50 intracortical depths) to identify eigenvectors of cytoarchitectural differentiation.

#### Neurosynth meta-analytic maps of functional diversity

Neurosynth^107^ collates data from over 15,000 previously published task-based functional MRI studies to facilitate coordinate-based meta-analyses of specific psychological terms. Meta-analytic maps derived from Neurosynth can be used to functionally decode groups of brain regions in terms of the mental processes that they support. Although Neurosynth data are not derived exclusively from studies conducted during youth, the psychological functions (e.g., vision, motor control, language, working memory, reward processing) supported by different cortical regions are broadly stable across adolescence and adulthood. We used the Neuroimaging Meta-Analysis Research Environment (NiMARE)^108^ to compute meta-analytic activation maps from version 0.7 of the Neurosynth database. In accordance with prior work on functional decoding^109,110^, we created meta-analytic maps for 123 psychological terms that were both available in Neurosynth and included in the Cognitive Atlas^111^, an ontology of cognitive neuroscience concepts. Term-specific meta-analytic activation maps were computed with NiMARE in volumetric space using the multilevel kernel density Chi-square analysis. Values in these meta-analytic maps are association test *z* values that quantify the degree to which activation in a given cortical location occurred more consistently in prior task-based functional MRI studies that mentioned a given term as compared to those that did not. Term-specific meta-analytic activation maps were mapped to the fslr 32k cortical surface, parcellated with the HCP-MMP atlas, and z-scored across cortical regions.

### Anatomical characterization

To determine whether regional variation in cortical R1 reflected variation in myelin density present in independent maps of cortical myeloarchitecture, we first averaged R1 across all 7 intracortical depths within each HCP-MMP region. We then computed the correlation between the resulting R1 map and the principal component of spatial variation across three independent myelin-sensitive measures. These measures included the T1w/T2w ratio obtained from contrast-based structural imaging^12^, magnetization transfer saturation derived from a multi-parameter mapping sequence^41^, and myelin basic protein gene expression^52^ as measured in transcriptomic data. The first principal component (PC1) of regional variation in these measures was computed using PCA. A non-parametric Spearman’s rank correlation was used to assess the monotonic relationship between R1 and PC1. The significance of this correlation was assessed with a conservative spatial autocorrelation-preserving null (*p*_SA-null_) using the BrainSMASH software version 0.11.0^112^.

BrainSMASH facilitates generative null modeling by producing surrogate (synthetic) brain maps with spatial autocorrelation that matches the degree of autocorrelation present in an empirical brain map. We used BrainSMASH to generate 1,000 surrogate maps that matched the spatial autocorrelation of the empirical R1 map. Surrogate maps were then correlated to PC1 to generate a null distribution of Spearman’s correlation values that was compared to the empirical R1 correlation to calculate *p*_SA-null_.

In addition to characterizing regional differences in average cortical R1, we surveyed regional differences in depth-dependent R1 profiles and examined whether these profiles paralleled those seen in high-resolution R1 data collected in 0.5 mm isotropic voxels at 7T. Publicly available high-resolution (0.5×0.5×0.5 mm) 7T R1 data collected from 10 healthy young adults (mean age = 22.6 ± 4.6 years) on a Siemens Terra by Cabalo et al.^113^ were used. As in the developmental dataset, R1 was derived in the high-resolution dataset from an MP2RAGE acquisition acquired with the following parameters: inversion times of 1000 ms (INV1) and 3200 ms (INV2), repetition time of 5170 ms, echo time of 2.44 ms, flip angles of 4 degrees, iPAT = 3. Residual B_1_ ^+^ transmit field inhomogeneity was addressed in R1 data with UNICORT. UNICORT-corrected R1 was sampled at 14 intracortical depths between pial and white matter boundaries based on surfaces generated using FastSurfer; the use of 14 depths facilitates comparisons between these high-resolution data (14 intracortical depths studied within 0.5 mm isotropic voxels) and the developmental data (7 intracortical depths studied within 1 mm isotropic voxels). Qualitative and quantitative analyses were employed to compare depth-dependent R1 profiles in the cortical ribbon between datasets.

Qualitative analysis focused on examining the shape of depth-dependent R1 profiles in different brain regions^82^. Quantitative analysis compared the skewness of regional depth-dependent profiles between developmental and high-resolution datasets, given known cortical heterogeneity in this property based on regional architectural complexity^11,82^. Skewness, or the third standardized moment of a probability distribution, is a statistical measure of the asymmetry of the distribution of a variable around its mean. In the present context, skewness provides insight into the distribution of R1 across superficial and deep cortical depths. Positive skewness (i.e., a right-skewed distribution) reflects relatively uniform levels of R1 across superficial and middle depths with a pronounced increase in deep depths. Negative skewness (i.e., a left-skewed distribution) indexes disproportionately low R1 in superficial as compared to all other depths. Skewness values near zero denote a distribution of R1 values that are symmetric around the mean, and thus uniformly gradual increases in R1 across depths. We calculated the skewness of depth-wise R1 in each cortical region in the developmental and high-resolution samples and compared these values across the brain between datasets using a Spearman’s correlation. Significance of the correlation was determined using BrainSMASH (*p*_SA-null_) based on 1,000 autocorrelation-preserving surrogates of the developmental skewness map.

### Superficial, middle, and deep cortex

We performed assessments to ensure that we could robustly differentiate R1 signal in superficial and deep portions of the frontal cortical ribbon. We first examined the thickness of the cortex in each frontal lobe HCP-MMP region using all longitudinal UNI images; cortical thickness was computed by FreeSurfer’s longitudinal processing stream. The goal of this examination was to verify that all regions of the frontal cortex were > 2 mm thick (at minimum) in all participants at all study sessions, as this would ensure that each cortical location contained signal from at least 2 independent (superficial and deep) voxels. Next, we directly calculated the number of independent voxels that were embedded in the cortical ribbon at all frontal lobe vertices in data from all participants. This procedure provided insight into the number of unique cortical compartments that R1 signal could be measured in. To determine the number of independent voxels contained in the cortex at each vertex, we assigned every voxel in an R1 image a unique number and sampled voxel numbers to surface vertices at the same 11 intracortical depths used in the R1 analysis (spanning 0% to 100% of cortical thickness in 10% increments). We then calculated the number of unique voxel values present at every frontal lobe vertex across only the 7 analyzed intracortical depths.

The above assessments demonstrated that the vast majority of the frontal lobe contained R1 signal from 3 or more unique voxel compartments across the 7 analyzed intracortical depths in scans from all participants, regardless of age. We therefore set out to differentiate superficial, middle, and deep cortical ribbon compartments by assigning each of the 7 intracortical depths to one compartment. In order to determine the most suitable compartment to assign each depth to, we used a data-driven approach to delineate compartment boundaries at depths where the depth-dependent pattern of R1 change shifted. This approach was motivated by the fact that myelination patterns shift between layers 2/3, layer 4, and layers 5/6, producing transition points between microstructurally distinct laminar compartments. To identify transition points, we computed the absolute second derivative of R1 across frontal lobe intracortical depths. The absolute second derivative quantifies the degree of change in the slope of the R1 profile within the cortical ribbon. A sharp acceleration or deceleration in the slope with increasing cortical depth results in a peak in the absolute second derivative and is indicative of a transition point in the statistical properties of laminar R1. We calculated the absolute second derivative by fitting a generalized additive model with frontal lobe R1 as the independent variable and a smooth function for intracortical depth.

Importantly, we calculated second derivatives to identify frontal lobe compartment boundaries in both the high-resolution dataset (0.5 mm isotropic; 14 intracortical depths) and the developmental dataset (1 mm isotropic; 7 intracortical depths). We began with the high-resolution dataset given that sub-millimeter resolution scans have greater sensitivity to detect subtle changes in intracortical R1 profiles that emerge from cortical layers. We then repeated the analysis in the developmental dataset and examined agreement in compartment boundaries between the two datasets. Based on convergent results between datasets, we ultimately averaged R1 across depths 1-2 (superficial compartment; 20-30% of cortical thickness), depths 3-5 (middle compartment; 40-60% of thickness), and depths 6-7 (deep compartment; 70-80% of thickness) for developmental analyses.

### Developmental analyses

#### Developmental modeling

We used generalized additive mixed models (GAMMs; semiparametric additive models) with penalized splines to characterize developmental change in R1 throughout different regions and compartments of the cortical ribbon. All GAMMs were fit using the mgcv package in R^114^. For regional analyses, GAMMs were fit independently to data from each HCP-MMP region. GAMMs allow for the characterization of both linear and non-linear age-dependent relationships by modeling age fits as flexible smooth functions (i.e., age splines) derived from a linear combination of weighted basis functions. Developmental GAMMs were fit with compartment-specific R1 as the dependent variable and age as a smooth term with a random intercept per participant. The smooth term models the group-level developmental trajectory of R1 in superficial, middle, or deep cortex. Thin plate regression splines were used as the smooth term basis set and the restricted maximal likelihood approach was used for selecting smoothing parameters. To prevent overfitting of the age smooth function, a maximum basis complexity (k) of 4 was used and spline wiggiliness was penalized. The significance of the smooth term *F-*statistic represents the significance of the developmental effect. When examining the significance of developmental effects across multiple frontal lobe regions, the false discovery rate correction was used to correct *p*-values for multiple comparisons (designated as *p*_FDR_). In addition to the main developmental models, a GAMM with an age-by-compartment interaction term was fit for R1 averaged across the frontal lobe to test whether frontal cortex age splines significantly differed across superficial, middle, and deep cortex. This age-by-compartment interaction GAMM was fit with compartment-wise R1 as the dependent variable, smooth terms for age and compartment, a tensor product (continuous-by-continuous) interaction between age and compartment, and a random intercept per participant nested within compartment. The significance of the tensor product interaction designates whether developmental fits significantly differed across cortical compartments.

To parameterize differences in the magnitude and timing of development throughout the frontal cortical ribbon, we extracted quantitative information regarding R1 development from age splines. Specifically, we calculated the average rate of R1 increase and the age of R1 maturation from the first derivative of the fitted age spline. All derivative statistics were implemented using the gratia package in R^115^. The first derivative of the spline was calculated with finite differences at 200 equally spaced age intervals, providing age-specific rates of developmental change. The average first derivative was then computed to index the average rate of R1 change across the spline; a greater average derivate is indicative of greater overall developmental change in the age range studied. To determine the age of R1 maturation, we identified the youngest age at which the first derivative was no longer significantly different from 0 by obtaining a simultaneous 95% confidence interval for the first derivative. This represents the age at which the rate of developmental change in R1 can no longer be statistically differentiated from 0 (two-sided).

#### Alignment to hierarchical and cytoarchitectural axes

In addition to investigating divergence in R1 developmental trajectories between superficial, middle, and deep cortex within the frontal lobe, we sought to characterize variability in R1 development across frontal lobe regions within each of the 3 cortical compartments. We hypothesized that regional variability in R1 development would align with both the hierarchical S-A axis and the histology-based axis of cytoarchitectural variation, given prior evidence that cortical plasticity unfolds hierarchically in youth^3,5,9,12^ and decreases with progressive laminar elaboration^53^. To assess these hypotheses, we first used Spearman’s correlations to relate both the S-A axis and the cytoarchitectural axis to regional rates of R1 change (the average first derivative) within superficial, middle, and deep cortex, producing compartment-specific correlation values. We next used multiple linear regressions to simultaneously assess the relevance of the S-A axis and the cytoarchitectural axis to regional R1 development within superficial, middle, and deep cortex. More specifically, we fit 3 independent regressions for superficial, middle, and deep cortex, each with the S-A axis and the cytoarchitectural axis as joint predictors of the rate of R1 change in each frontal region.

#### Principal component of maturational variability

To understand how overarching modes of laminar myelin maturation varied across the frontal lobe, we performed a PCA to identify the principal spatial component of maturational variability in R1. The goal of this PCA was to derive a low-dimensional spatial representation of regional differences in the shape of compartment-wise R1 developmental trajectories. To extract information about the shape of R1 trajectories, we calculated the average curvature of each GAMM-derived age spline. Curvature provides a measure of how fast a curve is changing direction at a given point, and thus how much it deviates from a straight line. In the present context, higher average curvature values designate quadratic-plateauing splines whereas low average curvature values indicate that splines are closer to linear. We calculated the curvature of regional age splines derived from R1 in superficial, middle, and deep cortex and used these data as the input to the PCA.

We divided the first principal component output by this PCA into deciles and characterized decile-specific developmental and functional properties. In particular, we calculated the average curvature of age splines and the average age of R1 maturation within superficial, middle, and deep cortex for each of the 10 deciles. We additionally performed functional decoding with Neurosynth to identify the primary psychological functions subserved by regions falling within each decile. Functional decoding was accomplished by averaging meta-analytic map z-scores across all regions in a decile for each of the 123 psychological terms and identifying the 5 terms with the highest scores in each decile. This approach to functional decoding allows for the mapping of distinct mental concepts and behaviors to cortical territories with distinct profiles of R1 maturation.

#### Developmental sensitivity analyses

We conducted sensitivity analyses to ensure that our primary developmental findings applied across sexes and were not being driven by potential confounding factors, including individual differences in cortical thickness, cortical gyrification, structural data quality, and cortical partial volume effects. For each sensitivity analysis, the main compartment-specific developmental GAMMs were refit while including an additional linear covariate. As in the main analyses, sensitivity analysis GAMMs were used to determine the significance of the developmental effect (FDR corrected), to visualize R1 developmental trajectories, and to quantify the average rate of R1 change and the age of R1 maturation. The PCA based on compartment-wise developmental trajectories was also re-derived using the curvature of age splines produced by sensitivity analysis GAMMs.

The first sensitivity analysis included sex as a covariate in GAMMs. Including sex as a covariate allowed us to examine both whether R1 significantly different between sexes, and whether our main developmental findings differed when covarying for participant sex. In addition to models that included a main effect of sex, we fit GAMMs with an age-by-sex interaction term, modeled by an ordered factor-smooth interaction, to test whether R1 developmental splines significantly differed between individuals assigned male and female at birth. The second sensitivity analysis included regional cortical thickness as a covariate to ensure that our R1 results were not attributable to age-related decreases in cortical thickness. The third sensitivity analysis included regional mean curvature as a covariate, which indexes the magnitude of folding present on the cortical surface and thus summarizes gyral and sulcal patterns^116^. This analysis aimed to mitigate any potential effects of inter-individual differences in curvature on our findings, given prior findings that R1 and local curvature are correlated at middle cortical depths^46^. Cortical thickness and mean curvature were both calculated as part of the FreeSurfer longitudinal processing stream and averaged in each HCP-MMP region for each longitudinal imaging session. The fourth sensitivity analysis included FreeSurfer’s euler number (averaged across hemispheres) as a covariate to rule out the possibility that age-dependent differences in the quality of cortical surface reconstruction explained our results. The euler number summarizes the topological complexity of reconstructed surfaces (calculated from the number of surface holes) and provides a reliable, quantitative metric of structural image quality^117^. The fifth and final sensitivity analysis included depth-specific measures of the cortex volume fraction in each region as a covariate. This analysis was undertaken to ensure that developmental findings were not impacted by partial volume effects.

### Associations with EEG aperiodic activity

EEG data collected from the same accelerated longitudinal sample of participants were integrated with cortical R1 measures to provide insight into associations between cortical myelin content and local circuit activity. We analyzed four minutes of eyes-open resting state EEG and used these data to isolate the aperiodic component of EEG activity. Aperiodic activity represents “background”, population neural activity that occurs independently of band-specific oscillations.

Following EEG preprocessing and quality control of source localized aperiodic activity (see below), both EEG and R1 data were available from 127 participants (190 matched longitudinal sessions) of the primary study sample of N = 140.

#### EEG preprocessing and source estimation

High-impedance EEG data were acquired using a Biosemi ActiveTwo 64-channel EEG system with a custom electrode cap configuration. Data were resampled to 150 Hz and preprocessed using a revised EEGLAB (version 2022.1^118^) processing pipeline that included the removal of flatline channels, low frequency drifts, noisy channels, short spontaneous bursts, and incomplete segments of data. Following initial preprocessing, independent component analysis was performed to identify eye-blink artifacts and remove their contribution to the signal. Additional details on EEG acquisition and preprocessing are available in McKeon et al.^60^.

Following preprocessing, we used Brainstorm version 03^119^ to generate source-localized EEG time series within the frontal cortex. Source localization uses each individual’s cortical and scalp surface reconstructions obtained from structural MRI to estimate where in the cortex the neural activity accounting for scalp recordings arose from. To perform cortical source estimation for each longitudinal EEG session, we used each participant’s corresponding session-specific surface anatomies computed with longitudinal FreeSurfer. The 64 EEG channels were first aligned to each participant’s scalp surface using automated head-point refinement and channel projection. This step positions EEG electrodes on each individual’s head based on their specific anatomy, standard fiducial landmarks, and manufacturer-provided sensor coordinates. Next, the forward solution was estimated using a realistic three-layer head model with the OpenMEEG^120^ Boundary Element Method (BEM) using downsampled FreeSurfer outputs. The three layers included brain, skull, and scalp with relative conductivities of 1, 0.0125, and 1, respectively. The forward solution calculates a lead field matrix that mathematically describes the physical process of neuronal current propagation from sources within the brain to each EEG electrode. Finally, the inverse solution was computed with the cortical surface as the source space using minimum-norm estimation with sLORETA normalization^121^. This process resulted in EEG time series at all vertices in individual-specific cortical ribbons. Time series were averaged across vertices within each HCP-MMP atlas region within the frontal lobe.

In order to estimate the aperiodic exponent from regional EEG time series, a power spectral density (PSD) was calculated in each region using Welch’s method (2 second Hamming window; 50% overlap). The FOOOF (specparam) algorithm was then implemented within Brainstorm to separate the aperiodic component from the periodic component (oscillatory peaks) of the PSD. The aperiodic component was extracted from the 1-50 Hz frequency range of each PSD and the exponent (i.e., the slope of the aperiodic component) was computed. FOOOF was implemented with the following parameters: aperiodic model = fixed, peak width limits = [0.5, 12], minimum peak height = 0, max number of peaks = 4. To ensure that we only analyzed EEG sessions with high quality aperiodic estimates and source reconstructions, we quantified the correspondence between session-specific and group-level maps of the frontal cortex aperiodic exponent. To enable this quantification, we first created a group-averaged map of the aperiodic exponent in all HCP-MMP frontal lobe regions. We then calculated the Pearson’s correlation between the group-average map and each individual-specific map and excluded any EEG sessions where the maps were anti-correlated. This resulted in the removal of 9 out of an original 199 EEG sessions and ensured that only EEG data with valid spatial patterns of source localized aperiodic activity were analyzed.

#### Modeling R1 associations with EEG

We used GAMMs to study age-related change in the EEG-derived aperiodic exponent, following the exact model specification described for R1 development GAMMs above. In addition, we extended R1 development GAMMs to study relationships between intracortical R1 and the aperiodic exponent in each frontal region while controlling for age. These GAMMs simultaneously tested 1) whether the aperiodic exponent was significantly associated with R1 and 2) whether the strength of this association differed between superficial and deep cortical compartments. Models thus included superficial and deep compartment R1 as the dependent variable, a smooth term that modeled the association between the exponent and R1 in deep cortex (main effect), and an ordered factor-smooth interaction between the exponent and compartment that indicated whether the R1-exponent relationship differed between superficial and deep cortex. Models additionally had a smooth term for age, a linear covariate for compartment, and a random intercept per participant. We set the maximum basis complexity of the aperiodic smooth term to 3 given that we did not expect R1-aperiodic relationships to largely deviate from linearity. As in the developmental analyses, GAMMs were fit separately for each frontal region and the false discovery rate correction was applied to correct main effect and interaction effect *p*-values across all regions (*p*_FDR_).

### Associations with cognitive performance

In addition to participating in MRI and EEG, study participants completed a set of developmentally-sensitive cognitive tasks. Given our prediction that myelination impacts learning and processing efficiency during prefrontal cortex-dependent tasks, we primarily studied performance on a two-stage sequential decision-making task^67,68^. This task was used to derive estimates of learning rates and processing speed. We additionally analyzed data from two eye movement (i.e., saccade) tasks, the anti-saccade and the visually-guided saccade, to test for differential relationships of prefrontal myelin with cognitive versus visuo-motor processing speed. Cognitive data were available from 131 participants (195 matched longitudinal sessions) of the original N = 140 for the two-stage sequential decision-making task and for 125 participants (187 matched longitudinal sessions) for the two saccade tasks.

#### Cognitive tasks

The two-stage sequential decision-making task^67^ asks participants to make a series of choices to earn rewards while they learn about stochastically changing reward contingencies. Specifically, participants make a binary choice in each stage of a two-stage trial and probabilistically receive a reward in the second stage. Participants aim to maximize rewards over 200 task trials delivered in 3 blocks. The first stage of each trial is characterized by fixed action-outcome relationships and the second stage by dynamically changing action-reward probabilities. In our developmentally-validated version of the sequential decision-making task^68^, individuals try to maximize “space treasure” rewards by first choosing between two spaceships to visit different planets (stage 1) and next choosing between two aliens on a planet (stage 2). More explicitly, during stage 1 of each trial, individuals choose between one of two spaceships. Both spaceships can take them to the same two planets, but they do so with differing probabilities. Spaceship 1 has a 70% chance of traveling to planet 1 and a 30% chance of traveling to planet 2; spaceship 2 has the inverse probability. These action-outcome probabilities are fixed (i.e., stable) throughout the task. During stage 2 of each trial, individuals choose between one of two aliens present on the planet they arrived at. They may or may not be rewarded with space treasure from the alien. The probability that they are rewarded by each alien slowly and independently drifts via a random walk over the course of the task. Critically, these shifting action- reward probabilities during stage 2 require continued learning of changing contingencies. Together, this two-stage design requires participants to build a stable understanding of the general task structure (stage 1) while dynamically updating their understanding of shifting reward probabilities (stage 2). Prior work has validated that individuals as young as 8 years old learn the task structure, the transition structure (i.e., which spaceship travels to which planet most frequently), and accrue rewards non-randomly^68^.

The anti-saccade and visually-guided saccade tasks ask participants to make eye movements away from (anti-saccade) or towards (visually-guided saccade) peripherally presented stimuli. These two tasks require, respectively, the intentional inhibition versus engagement of a reflexive, automatic saccade response. The acquisition and scoring of the anti-saccade^122^ and visually-guided saccade^60^ tasks are described extensively in published literature from our group. Participants completed 48 trials of the anti-saccade task during which eye tracking was performed with a long-range optics eye-tracking system from Applied Science Laboratories (Model 6000) with a sampling rate of 60 Hz.

During each of the 48 trials, a red fixation cross was presented at the center of a black screen for a variable time between 500 and 6000 ms, followed by a 200 ms black screen. A yellow cue dot then appeared for 1000 ms pseudo-randomly in one of four positions on the horizontal meridian (2%, 33%, 66%, or 98% of screen width). Participants were instructed to direct their gaze away from the yellow cue to its horizontal mirror location, i.e., to refrain from looking at the cue and instead saccade to the opposite side of the screen. The visually-guided saccade task was acquired independently of the anti-saccade over 3 runs of 2 trials (60 trials total). Each trial began with a fixation cross presented for 1000 ms, after which a peripheral cue was presented at an unknown location along the horizontal midline. Participants were instructed to saccade to the cue and maintain fixation until the cue disappeared. Eye tracking in the visually-guided saccade task used horizontal electrooculogram channels recorded from facial muscles that were calibrated to eye position along the screen’s horizontal meridian.

#### Learning rates

We used the two-stage sequential decision-making task to derive measures of learning rate.

Given that the two stages of the task are designed to require different levels of dynamic learning, we computed learning rates for stages 1 and 2 of the task separately. To operationalize learning rates, we used the hBayesDM package^123^ in R to fit a hierarchical Bayesian reinforcement learning model to trial-level task data from all participants. Reinforcement learning models are computational models that parameterize latent neurocognitive processes based on participants’ decisions in a task.

Hierarchical Bayesian analysis, which estimates both group-level and individual-level parameter estimates (e.g., learning rates) simultaneously, is considered the gold-standard approach to model parameterization^123–126^. The hierarchical Bayesian approach uses group-level population distributions of parameters as priors when estimating individual-level parameters, which produces more robust and stable estimates of model parameters per person, thereby enabling better individual inference^123–125^. Posterior inference in the hBayesDM model was conducted using the No-U-Turn Sampler (NUTS), which is a part of the Hamiltonian Monte Carlo framework for Markov chain Monte Carlo (MCMC) sampling. The model was fit with 4 chains, each running for a total of 4,000 iterations, which included 1,000 warmup samples for each chain. Hyperparameters consisted of a population-level mean and standard deviation. Individual-level parameters were assigned weakly informative priors, which help regularize the estimates and ensure that the individual-level parameters are not overly constrained.

The hBayesDM reinforcement learning model analyzed in this work is a 7-parameter model fit identically as in our prior study that used the same sequential decision-making task in a similar adolescent and young adult sample^127^. The model comes pre-programmed with the hBayesDM package based on the widely adopted reinforcement learning model originally applied by Daw et al.^67^ to this same task. The 7 model parameters include stage 1 and stage 2 learning rates (α_1_ and α_2_), stage 1 and stage 2 inverse temperature (β_1_ and β_2_), a perseverance parameter (π), a model-based weight (w), and an eligibility trace (λ). These parameters have been described extensively in prior studies^67,128^. The learning rate parameter analyzed here captures how quickly an individual updates their expectations and choices based on new information (namely, recent outcomes in the task). Higher learning rates are indicative of greater sensitivity to more recent information and faster updating, and thus a faster step size or speed of learning^126^. Of note, after fitting the 7-parameter model, we confirmed that it had better performance than alternative model formulations with 6 parameters only (identical to the 7-parameter model, but without the eligibility trace) and 4 parameters only (across-stage learning rate, across-stage inverse temperature, perseverance, and model-based weight). Better performance was indicated by a lower Leave-One-Out Information Criterion (LOOIC) and Widely Applicable Information Criterion (WAIC).

#### Processing speed

Response times during cognitive tasks were used as readouts of processing speed. Given that the two-stage sequential decision-making task and the anti-saccade task require decision making and response inhibition respectively, response times on these tasks index the speed of processing when endogenously employing higher-order executive functions. In contrast, the visually-guided saccade task only requires simple, reflexive foveation of a stimulus; response times on this task thus reflect exogenously-driven sensorimotor processing. For the sequential decision-making task, average response times were calculated for stage 1 and stage 2 of the task independently as each stage requires making a decision based on different information^68^. Response time on this task is the time it takes for participants to make a binary choice through a keyboard press following presentation of task stimuli.

For the anti-saccade task, response time was calculated as the average latency to initiate a saccade on correct trials of the task. We only considered correct trials so as to only analyze timepoints where higher-order inhibitory control over reflexive responding was engaged successfully. Correct trials are defined as those in which the first eye movement during the saccade epoch with a velocity greater than or equal to 30◦/sec is made to the mirror location (opposite in direction) from the peripheral cue. Consistent with established protocol, we excluded trials with eye blinks as well as trials with response latencies less than 100 ms (express saccades) or greater than 1000 ms (non-response). For the visually-guided saccade task, response time was calculated as the average latency to initiate a saccade on all trials of the task while again excluding trials with response latencies less than 100 ms or greater than 1000 ms as well as trials with position errors greater than 23 degrees.

#### Developmental change in cognitive measures

We studied whether stage-specific learning rates and processing speed on the sequential decision-making task significantly changed with age in both the full sample using GAMMs and in participants with longitudinal data using person-mean centered regression. In the full sample, we adopted the model specification described for R1 development GAMMs above: models included the cognitive measure as the dependent variable, a smooth term for age, and random intercepts per participant. To evaluate whether within-person longitudinal change in cognitive measures paralleled GAMM results obtained at the group-level, we mean centered each participant’s age at each longitudinal timepoint. We then included person-mean centered age as a predictor of each cognitive measure in linear mixed effects models with random intercepts per participant. These models isolate within-person effects of age on task performance.

#### Modeling R1 associations with cognition

The primary set of cognitive analyses examined stage-specific learning rates and processing speeds from the two-stage sequential decision-making task. To first offer insight into whether learning rates and processing speeds systematically differed between the two stages of the task, we implemented two-sided paired samples t-tests. For analyses relating sequential decision-making task measures to cortical R1, we followed the same modeling procedure described above for EEG. Specifically, we fit region-specific GAMMs with superficial and deep compartment R1 as the dependent variable, the cognitive measure as a smooth term, an ordered factor-smooth interaction between the cognitive measure and cortical compartment (superficial versus deep), a smooth for age, a linear covariate for compartment, and random intercepts. As above, in these models the base cognitive term is a main effect modeling the association between task measures and R1 in deep cortex, whereas the interaction term tests for a difference in the nature of this association in superficial cortex.

Interaction models provided no evidence for significant differences in the strength of the relationship between sequential decision-making task measures and R1 in superficial and deep cortex. We therefore amended GAMMs to fit final models with a global cognitive smooth that captured the relationship between sequential decision-making task measures and R1 in both deep and superficial cortex. These GAMMs were used for statistical inference and were fit identically as above while removing the ordered factor-smooth interaction term. We adopted the same model formulation to study anti-saccade and visually-guided saccade response times in cognitive specificity analyses.

For all cognition analyses, we corrected for multiple comparisons across frontal lobe regions using the false discovery rate correction (*p*_FDR_).

## Data availability

Raw neuroimaging data from this study are available upon request. This study incorporated previously published cortical maps, including the S-A axis (accessed in fslr 32k space from https://github.com/PennLINC/S-A_ArchetypalAxis), T1w/T2w ratio developmental data (available for download from https://balsa.wustl.edu/study/P2DmK), left hemisphere myelin basic protein gene expression values (released in fslr 32k space through https://github.com/kwagstyl/MAGICC), and a histological axis of cytoarchitectural variation (Hist-G2 obtainable in fslr 32k space from https://bigbrainwarp.readthedocs.io/en/latest/pages/toolbox_contents.html). It additionally used Neurosynth version 0.7 data, released on github (https://github.com/neurosynth/neurosynth-data) and volumetric and surface templates made available through TemplateFlow (https://www.templateflow.org/).

## Code availability

All original code generated for this study is available on github at https://github.com/LabNeuroCogDevel/corticalmyelin_maturation. In addition, a detailed guide to code implementation describing all analytic steps and statistical analyses is provided along with the github repository at https://labneurocogdevel.github.io/corticalmyelin_maturation. Original study code integrated publicly available software. MRI processing utilized a containerized version of the SPM BIDS app (available at https://hub.docker.com/r/bids/spm/), a containerized FreeSurfer BIDS app (available at https://hub.docker.com/r/bids/freesurfer; tag freesurfer:7.4.1-202309), a Neuromaps container (available at https://hub.docker.com/r/netneurolab/neuromaps; tag 0.0.4), AFNI (available at https://afni.nimh.nih.gov/pub/dist/doc/htmldoc/index.html; version 23.1.10), and Connectome workbench (available at https://github.com/Washington-University/workbench/releases; version 1.5.0). EEG processing used EEGLab (available at https://eeglab.org/download/; version 2022.1), FOOOF (available at https://fooof-tools.github.io/fooof/; version 1.0.0), and Brainstorm 03 (available at https://github.com/brainstorm-tools/brainstorm3). Neurosynth analyses relied on NIMARE (available at https://nimare.readthedocs.io/en/latest/installation.html). Spatial autocorrelation preserving nulls were generated with BrainSMASH version 0.11.0 (https://brainsmash.readthedocs.io/en/latest/index.html). Reinforcement learning models were fit with hierarchical Bayesian modeling with the model specified under “Two-Step Task – Hybrid Model, with 7 parameters (original model)” in hBayesDM (https://hbayesdm.readthedocs.io/en/develop/models.html). All statistical analyses were conducted in R version 4.2.3 (https://cran.r-project.org/bin/macosx/).

## Acknowledgements

This work was supported by the National Institutes of Health (NIH), including T32MH016804 and T32MH018951 to VJS, T32AA007453 to DP, F31MH132246 to SDM, and R01MH067924 to BL. Additional funding was provided by the Staunton Farm Foundation. This work used Bridges-2 at the Pittsburgh Supercomputing Center through allocation soc23000rp from the Advanced Cyberinfrastructure Coordination Ecosystem: Services & Support (ACCESS) program, which is supported by National Science Foundation grants #2138259, #2138286, #2138307, #2137603, and #2138296.

We are grateful to personnel at the Magnetic Resonance Research Center (MRRC) at UPMC Presbyterian, especially Chan Moon and Hoby Hetherington, for their valuable assistance in planning, implementing, and performing the 7T imaging acquisitions. We also thank the University of Pittsburgh Clinical and Translational Science Institute (CTSI) for their support in recruiting participants, as well as their support by the NIH through UL1TR001857.

## Author Contributions

VJS designed the study with support from FJC and BL. AF acquired the neuroimaging data under the supervision of FJC and BL and using resources acquired by BL. VJS curated and quality controlled the MRI data with support from WF. VJS developed and implemented the MP2RAGE processing pipeline. SDM processed the EEG data. DP analyzed the two-stage sequential decision-making task data. SDM and WF analyzed data from the two saccade tasks. VJS carried out all statistical analyses with guidance from FJC. VJS wrote all study-specific code and created all figures. VJS wrote the original manuscript. All authors reviewed and revised the manuscript. FJC and BL provided scientific mentorship throughout the lifecycle of the project.

## Competing Interests

The authors declare no competing interests.

## Notes

### Competing Interest Statement

The authors have declared no competing interest.

### Summary of Updates

This revision addresses the differentiation of myelin-related neuroimaging signal between cortical layers through the inclusion of a high-resolution (0.5 mm isotropic) 7T R1 dataset and an updated methodological approach for distinguishing R1 signal in superficial, middle, and deep cortex.

