## Supplementary Figures for "Heterochronous laminar myelination in the human prefrontal cortex"

**Affiliations**

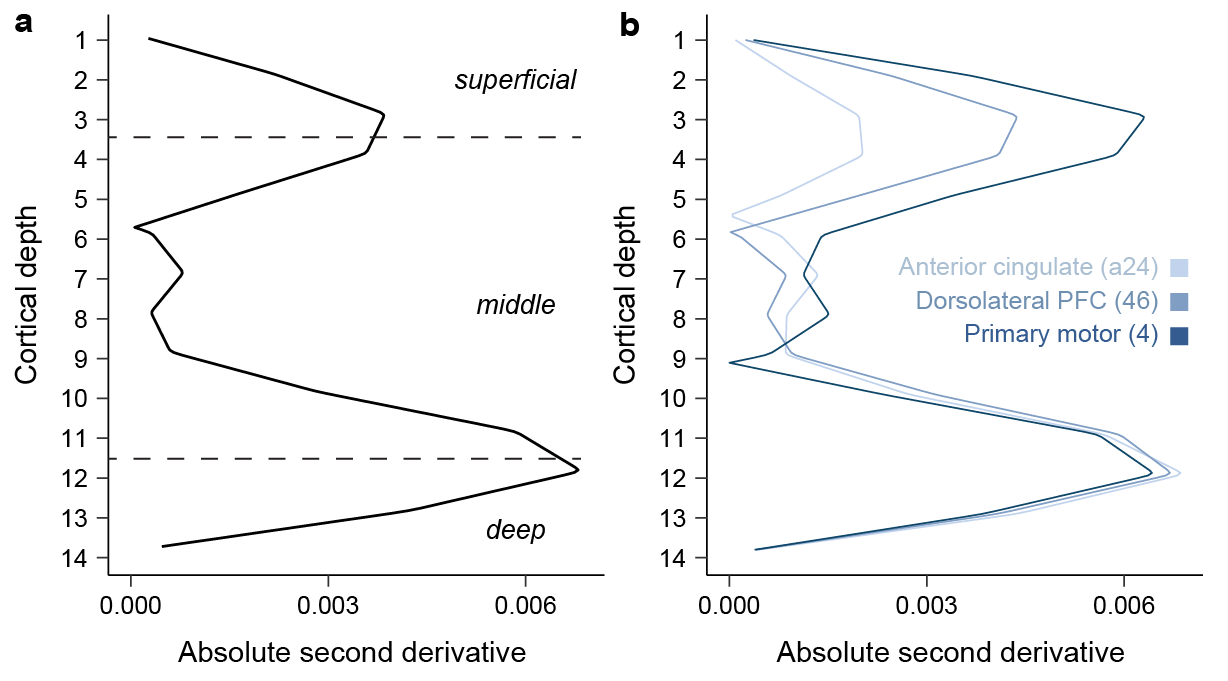

**Supplementary Figure 2.1. Differentiating R1 in superficial, middle, and deep compartments of the frontal cortex in high-resolution data.** The absolute second derivative was used to identify where in the cortical ribbon depth-dependent change in R1 robustly accelerated or decelerated, suggestive of layer-related shifts in myelination properties. The derivative analysis was applied to frontal lobe R1 data sampled at 14 intracortical depths in the high-resolution dataset, leveraging the capacity for 0.5 mm isotropic resolution data to better resolve cortical layer properties. Peaks in the absolute second derivative—indicative of transition points in the laminar R1 profile—were identified near cortical depths 3-4 and 11-12 both when R1 was averaged across the frontal lobe (**a**) and examined within individual cortical regions with distinct laminar architectures (**b**).

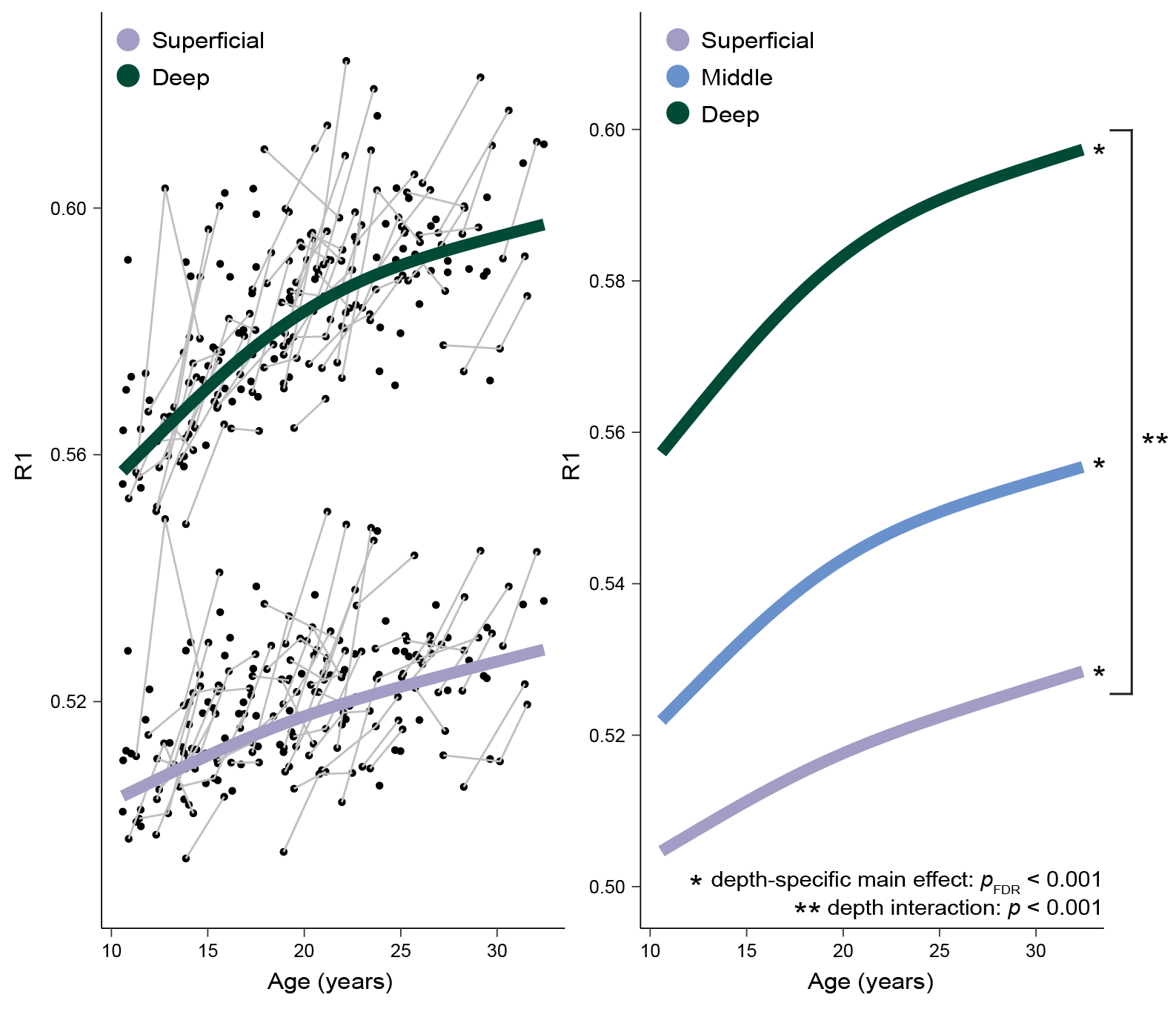

**Supplementary Figure 3.1. R1 developmental trajectories vary within the frontal cortical ribbon.** Developmental trajectories of frontal cortex R1 diverge between superficial, middle, and deep cortex. Compartment-specific developmental trajectories, derived from GAMM smooth functions, are plotted simultaneously for superficial and deep cortex overlaid on participant-level data (left) and for superficial, middle, and deep cortex (right). The R1 developmental trajectory is shallow yet protracted in superficial cortex and steep yet plateauing in mid-to-deep cortex.

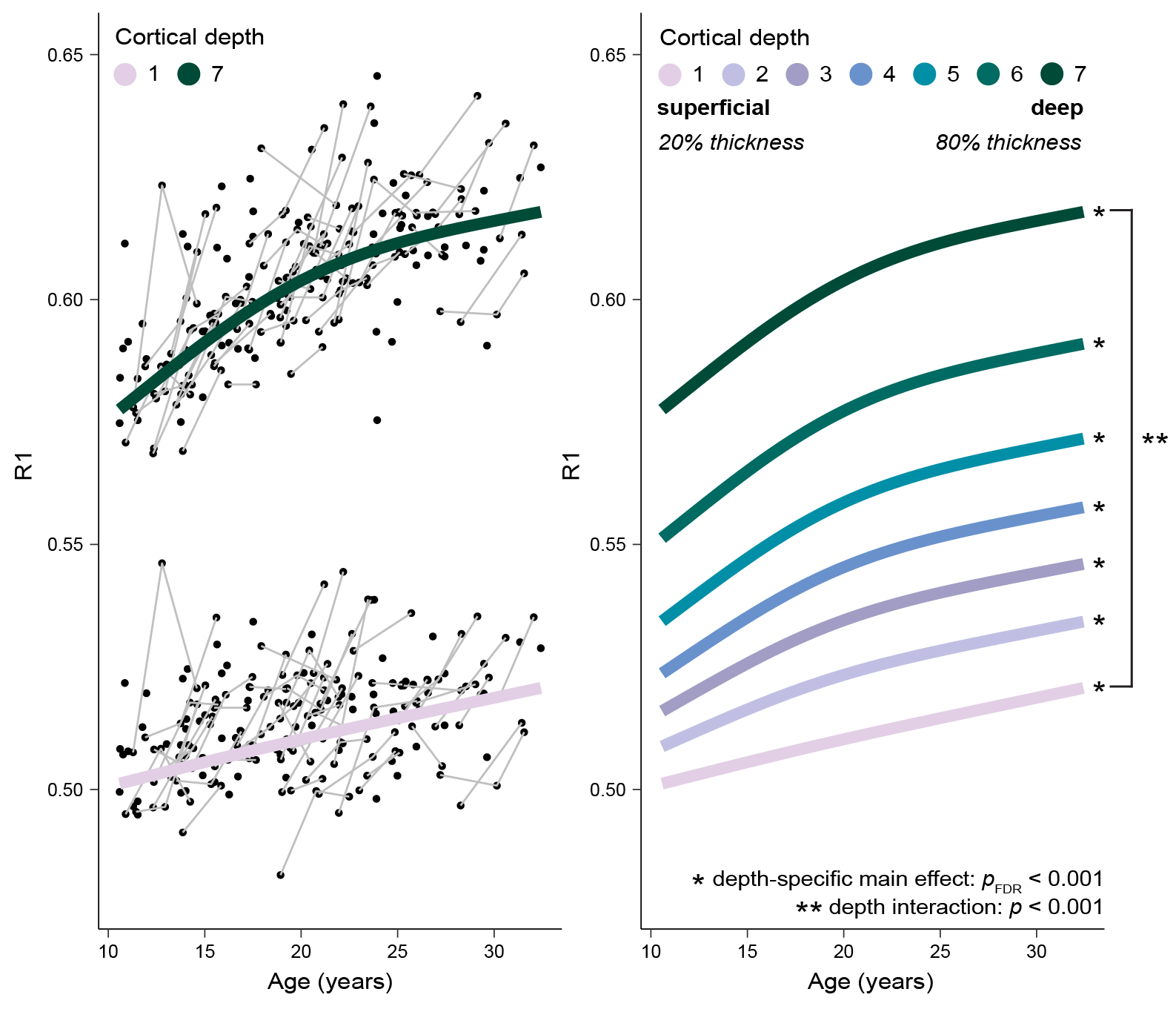

**Supplementary Figure 3.2. R1 developmental trajectories diverge between superficial and deep cortex across individually sampled intracortical depths.** Developmental trajectories of frontal cortex R1 differ between relatively more superficial and deep cortex when defining trajectories at 7 intracortical depths with minimal partial voluming. Depth-specific developmental trajectories (GAMM smooths) are plotted for the most superficial and deep depths overlaid on participant-level data (left) and for 7 intracortical depths ranging from 20% to 80% of cortical thickness in 10% thickness increments (right).

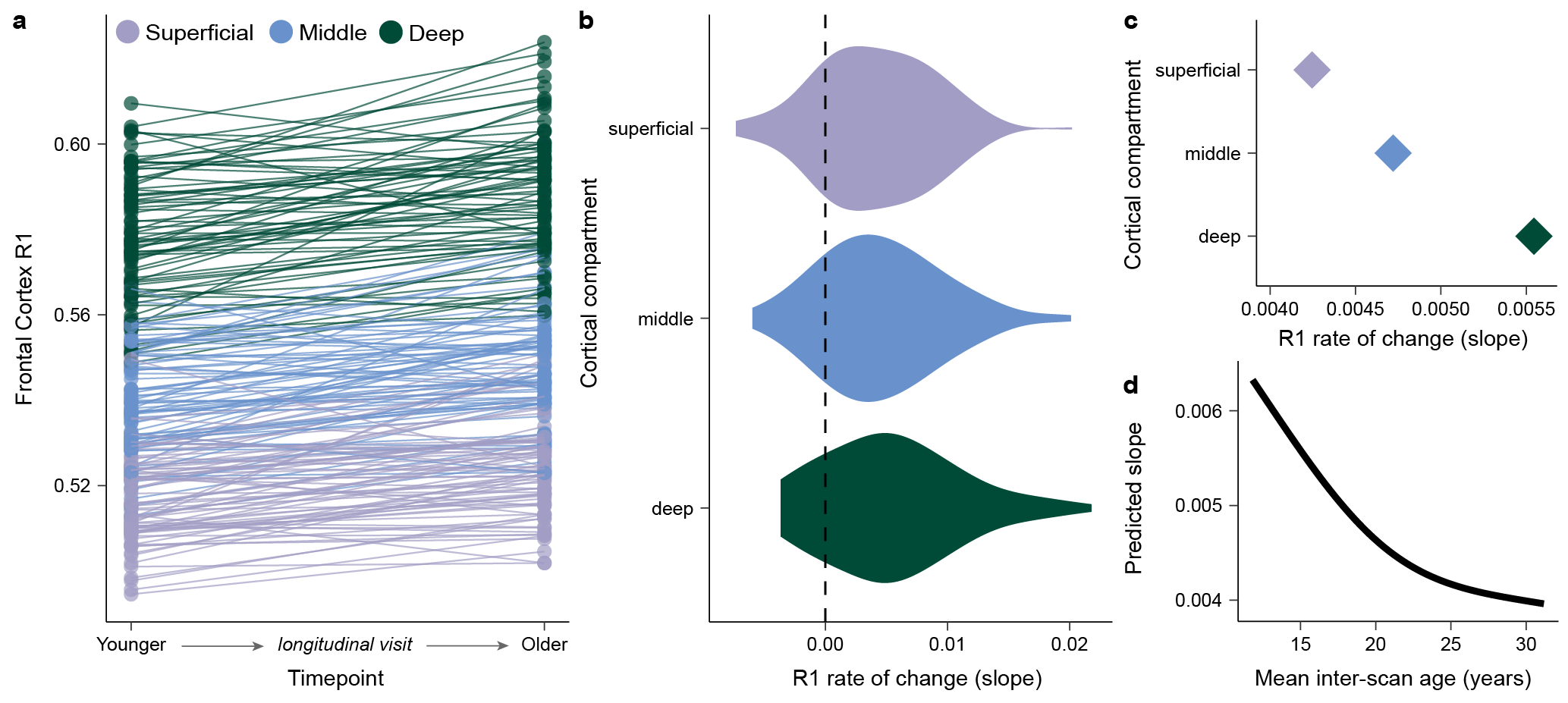

**Supplementary Figure 3.3. Convergent findings from within-person estimates of R1 longitudinal development.** Within-person change in frontal cortex R1 provides convergent evidence for age-related increases that grow in magnitude with greater distance into cortex and decelerate at the end of adolescence. **a**) Within-person change in frontal cortex R1 between longitudinal imaging timepoints is visualized for superficial, middle, and deep cortex for all participants with longitudinal data. Lines connect longitudinal imaging sessions from the same participant; both session 1 to session 2 paired timepoints and session 2 to session 3 paired timepoints are included. Ubiquitous within-person increases in frontal R1 are evident. **b**). Within-person quantification of longitudinal R1 change confirms robust age-related increases in the majority of participants in superficial, middle, and deep compartments of the frontal cortical ribbon. The longitudinal rate of R1 change between paired imaging timepoints was quantified for every participant by calculating Δ R1/Δ age, deriving person-specific age slopes. Plotting this age slope for all participants in superficial, middle, and deep cortex confirmed that the rate of frontal R1 change was positive (> 0) in the majority of participants in all 3 compartments. Specifically, the rate of R1 change was positive for 83% of longitudinal sessions in the superficial compartment, 83% in the middle compartment, and 80% in the deep compartment. **c**) Within-person estimates of R1 age slopes get larger, on average, when moving from superficial to deep cortex. The average value of the depth-specific distributions shown in **b** are plotted. **d**) Within-person R1 change is maximal in younger participants and decelerates with increasing participant age. A GAMM was used to model how the longitudinal rate of R1 change varied across the age range under study. The dependent variable of the GAMM was person-specific rates of change in all compartments (age slopes from **b**). Independent variables included a smooth function for participant mean inter-scan age (average age across the two imaging timepoints) and a random intercept per compartment. Plotting the GAMM-predicted rate of longitudinal R1 change across age (i.e., fitted values for the Δ R1/Δ age slope) revealed that rates of change were largest at the youngest ages and decelerated at older ages, providing evidence for a slowing of within-person developmental increases in R1 at the end of adolescence.

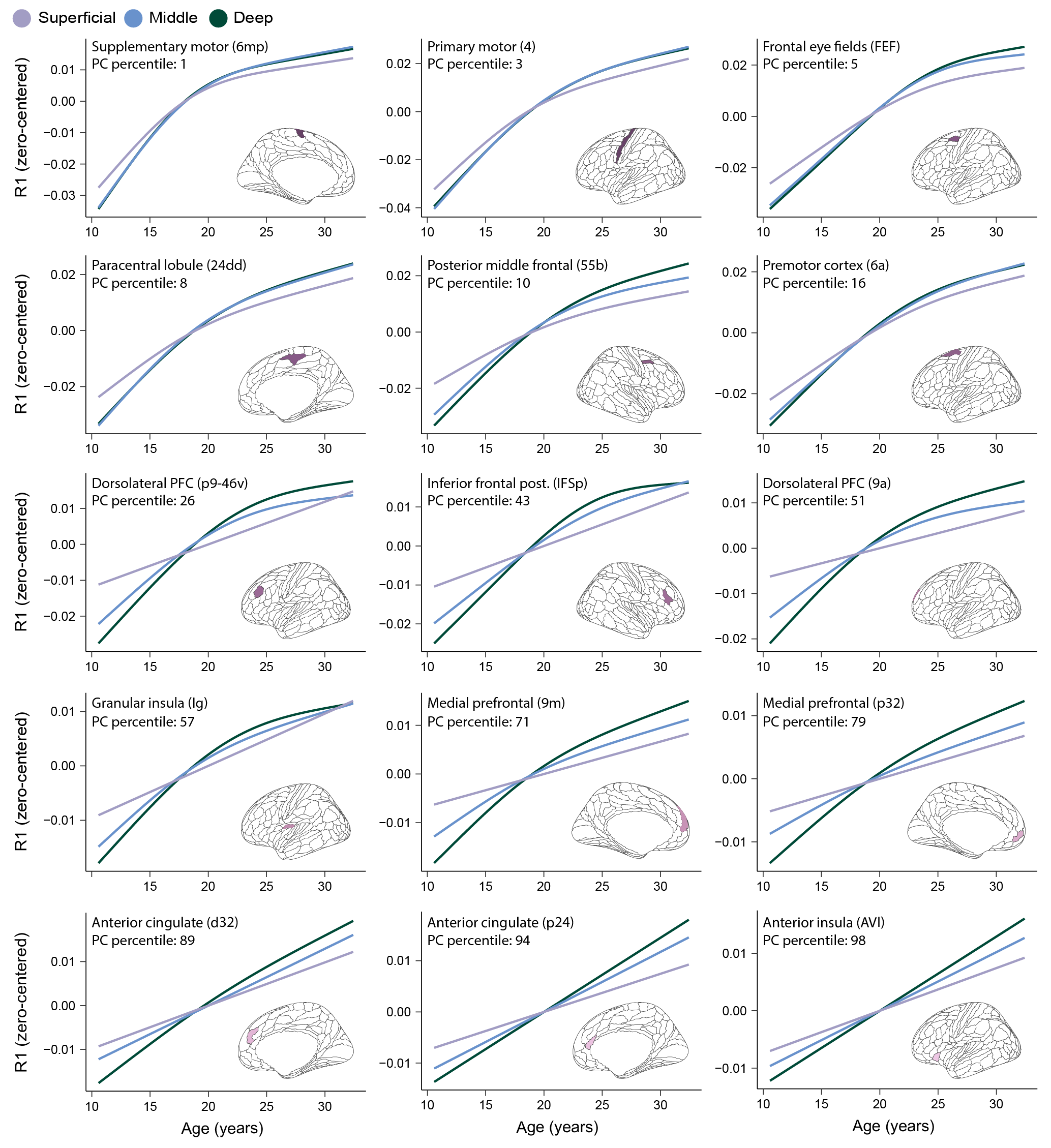

**Supplementary Figure 5.1. The principal component of maturational variability captures a spectrum of laminar R1 developmental profiles across the frontal lobe.** Developmental trajectories of R1 are shown for superficial, middle, and deep cortical compartments in frontal lobe regions that localized to different sections of the principal component of maturational variability. Frontal regions are ordered according to their percentile rank in the principal component (PC). Trajectories are zero-centered GAMM smooth estimates (i.e., splines) that reflect the partial effect of age on R1 accounting for model covariates.

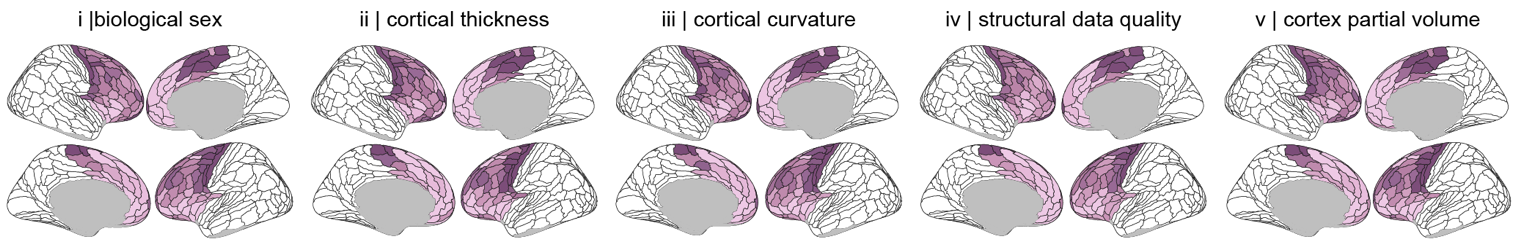

**Supplementary Figure 6.1. The principal component of maturational variability is stable across sensitivity analyses.** The principal component of laminar maturational variability, derived from a principal component analysis applied to regional R1 developmental trajectories, is highly stable across sensitivity analyses that covaried developmental models for i) biological sex, ii) cortical thickness, iii) cortical curvature, iv) structural data quality, and v) cortex partial volume effects.
